# Moss functional traits are important drivers for moss and underlying soil bacterial communities: evidence from a chronosequence in an Icelandic glacier forefield

**DOI:** 10.1101/2022.04.13.488162

**Authors:** Ingeborg J. Klarenberg, Christoph Keuschnig, Alejandro Salazar, Liane G. Benning, Oddur Vilhelmsson

## Abstract

Mosses are among the first colonizing organisms after glacier retreat and can develop into thick moss mats during later successional stages. They are key players in N_2_ fixation through their microbiome, which is an important process for nutrient build-up during primary succession. How these moss-microbe interactions develop during succession is not well-studied and is relevant in the light of climate change and increased glacier retreat.

We examined how the bacterial communities associated with two moss species of the genus *Racomitrium* and the underlying substrate, as well as moss traits and nitrogen fixation, develop along a successional gradient in the glacier forefield of Fláajökull in southeast Iceland. In addition, tested whether moss functional traits, such as total carbon (TC) and nitrogen contents (TN) are drivers of moss and underlying soil bacterial communities.

Although time since deglaciation did not affect TN and moisture content, TC and shoot length increased with time since deglaciation. Moss and underlying soil bacterial communities were distinct. While the soil bacterial community structure was driven by the time since deglaciation and moss C/N ratios, the moss bacterial community structure was linked to time since deglaciation and moss moisture content. Moss N_2_-fixation rates were linked to bacterial community composition and *nifH* gene abundance rather than moss TN or time since deglaciation. This was accompanied by a shift from autotrophic to heterotrophic diazotrophs.

Overall, our results suggest that there is little lateral transfer between moss and soil bacterial communities and that moss traits and time since deglaciation affect moss and soil bacterial community structure. In addition, moss N_2_-fixation rates are determined by bacterial community structure, rather than moss traits or time since deglaciation.

## 1. Introduction

Formerly ice-covered terrains are becoming increasingly exposed as glaciers retreat due to climate change (Roe, Baker, and Herla 2017). Such glacier forefields are subject to rapid ecosystem development, with microbial communities as the first colonizers. These early colonizing microbial communities are responsible for the first stages of soil development, which often involves the formation of a Biological Soil Crusts capable of stabilizing the soil and of fixing carbon (C) and nitrogen (N) (Bradley, Singarayer, and Anesio 2014; Breen and Lévesque 2008). The subsequent increase in C and N availability facilitates the colonization of other organisms, such as mosses (Vilmundardóttir, Gísladóttir, and Lal 2015b). Mosses can develop into thick moss mats during succession, especially in regions with high precipitation and little competition from higher plants (Tallis 1958). Moss establishment further enhances soil development in newly exposed terrain, by contributing to N (Arróniz-Crespo et al. 2014; Bowden 1991; Vilmundardóttir, Gísladóttir, and Lal 2015b), retaining moisture, and contributing to organic matter build-up (Wietrzyk-Pełka et al. 2020; Vilmundardóttir, Gísladóttir, and Lal 2015b), which additionally promotes soil microbial growth (Bardgett and Walker 2004). Thus, while microbial communities create the conditions necessary for plant establishment, plants influence microbial communities, for instance via litter inputs (Fanin, Hättenschwiler, and Fromin 2014). Despite an increasing number of studies linking the development of soil microbial communities to establishment of plants in glacier forefields (Arróniz-Crespo et al. 2014; Knelman et al. 2012; 2018; Bueno de Mesquita et al. 2017), we have a very limited understanding of the dynamics of moss-associated bacterial communities during ecosystem development in these environments.

Due to their diazotrophic microbiome (Ininbergs et al. 2011), mosses are the most important source for new N in Arctic ecosystems (Rousk, Sorensen, and Michelsen 2017). As glacier forefields are typically nutrient limited, moss microbiomes may be key players in biogeochemical N cycling during primary succession (Arróniz-Crespo et al. 2014). N_2_-fixation rates are variable and can be influenced by moss species (Stuart et al. 2020; Jean et al. 2020), N availability (Arróniz-Crespo et al. 2014), moisture (Rousk, Sorensen, and Michelsen 2018), temperature (Rousk 2017), diazotroph composition (Ininbergs et al. 2011), diazotroph abundance (Arróniz-Crespo et al. 2014) and diazotroph activity (Warshan et al. 2016) throughout succession.

Moss-associated bacterial community composition may also be driven by host identity (Holland-Moritz et al. 2018). Moss traits such as C and N content, may affect moss-associated bacterial community composition, similarly to how phyllosphere microbial taxa are linked to leaf traits (Yunshi Li et al. 2018; Laforest-Lapointe et al. 2017). These moss traits can change during succession. For instance, Sphagnum and bryophyte C/N ratio increased with peatland succession (Laine et al. 2021) and time since deglaciation in Chilean glacier forefields (Arróniz-Crespo et al. 2014). These changes might subsequently affect the composition of the moss-microbiome.

The development of a plant-microbiome during succession may furthermore depend on where the microbes are inherited from (Poosakkannu et al. 2017). Plant-associated microbial communities are thought to be mainly inherited from the surrounding soil which is also referred to as horizontal transfer (Compant et al. 2019). While mosses might not have a large rhizosphere, some have rhizoids and are thus connected to the soil. In higher plants, microorganisms can also be transferred vertically, via the seed (Hardoim et al. 2012). For mosses, microbial organisms might indeed be transferred between the sporophyte and the gametophyte (Bragina et al. 2012) or via vegetative regeneration (Tallis 1959). Depending on which source is more important for the composition of moss-associated bacterial communities, successional changes in soil microbial communities can be reflected in changes in the moss microbiome, or alternatively the moss microbiome may stay relatively stable throughout the development of a succession.

As moss cover increases the amount of organic carbon, moisture and nutrient content in soil (Bragazza et al. 2019; Breen and Lévesque 2008; Bardgett and Walker 2004), it may also indirectly influence the underlying soil bacterial communities (Juottonen et al. 2020).

Here we examine the bacterial communities of two moss species of the genus *Racomitrium* and the underlying substrate along a successional gradient in the glacier forefield of Fláajökull, in southeast Iceland. Mosses of the genus *Racomitrium* are important colonizers in Icelandic glacier forefields (Vilmundardóttir, Gísladóttir, and Lal 2015a; Glausen and Tanner 2019).

We hypothesized that: (i) moss total N (TN) and moss total C (TC) increase with time since deglaciation; (ii) changes in moss functional traits (such as TN and TC) and time since deglaciation lead to shifts in moss-associated bacterial communities and the underlying soil bacterial community. We also hypothesized that moss-associated N_2_-fixation rates and *nifH* gene abundance: (iii) will decrease with time since deglaciation as TN increases since high levels of N availability could reduce physiological needs to fix N; (iv) increase with moss moisture content and (v) depend on bacterial community composition.

## 1. Materials and methods

The chronosequence we studied lies in the pro-glacial area of Fláajökull glacier (64.328124°; -15.527791°), which is an outlet glacier on the south-eastern side of the Vatnajökull icecap (Figure 1). The Fláajökull glacier forefield is characterized by a number of moraine ridges and other landforms such as drumlins and eskers (Evans, Ewertowski, and Orton 2016; Jónsson et al. 2016). The oldest moraine dates from the glacier’s furthest advance towards the end of the Little Ice Age in 1894 (Hannesdóttir et al. 2015). The extent of the glacier in the last 120 years has been estimated using multiple dating methods, including glaciological methods, lichenometry and historical records (Evans, Ewertowski, and Orton 2016; Dabski 2002; Icelandic Glaciological Society 2018). Our furthest sampling point in 2018 lays more than 3000 m from the front of the glacier.

**Figure 1.**
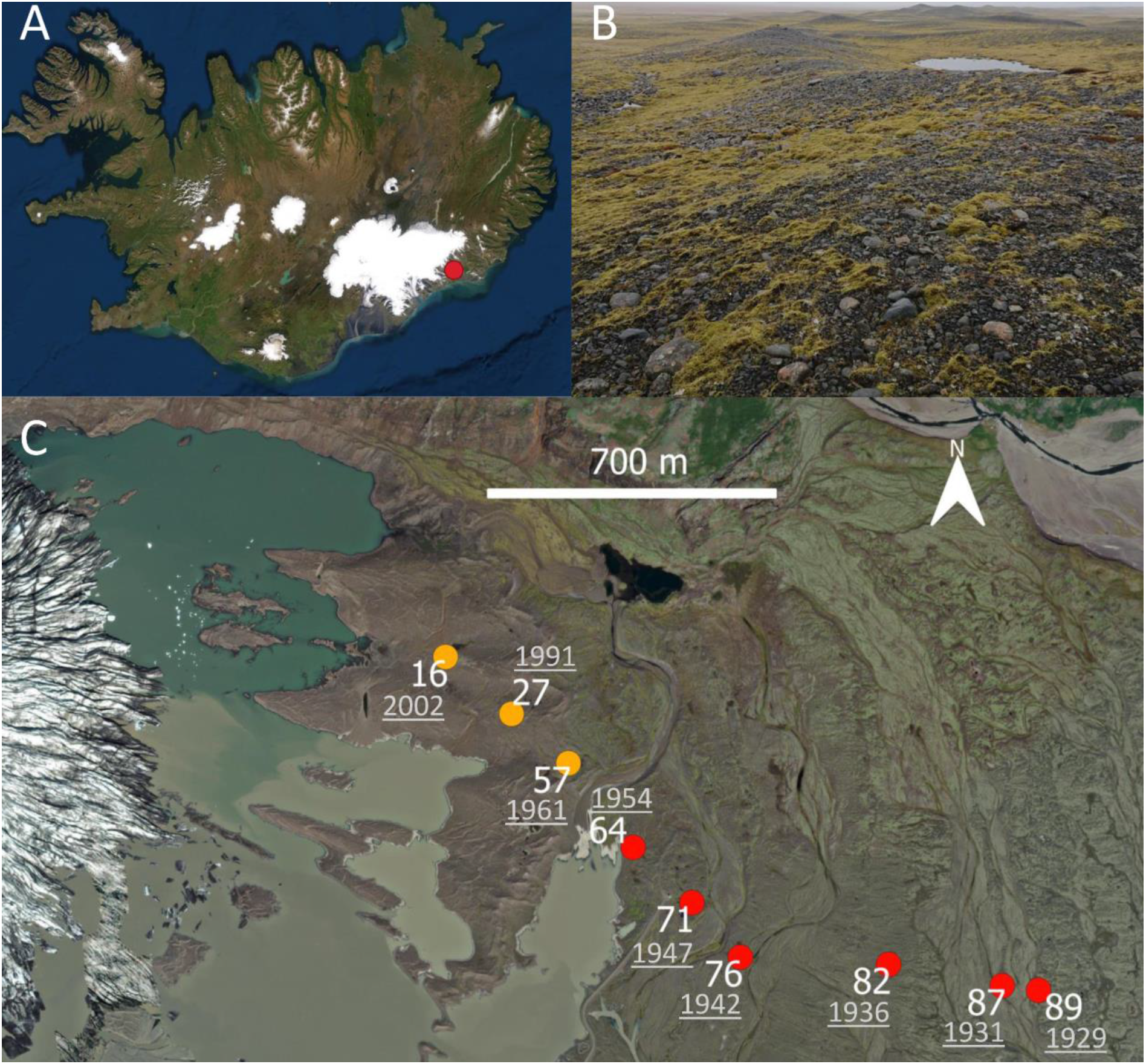
Overview of A) the location of the Fláajökull glacier forefield, B) *Racomitrium* spp. at the sampling site and C) the sampling locations along the chronosequence. The sites with *R. ericoides* are depicted in yellow and the sites with *R. lanuginosum* in red. At each sampling location the time since deglaciation in 2018 and year of deglaciation are indicated. Time since deglaciation was determined after Evans et al. 2016 and the Icelandic Glaciological Society, 2018.

The two closest weather stations are located in Fagurhólsmýri (72 km) and Höfn (18 km), which have a mean annual temperature of 4.8 °C and 4.6 °C respectively and a mean annual precipitation of 1814 mm and 1381 mm respectively. The climate can be described as sub-polar oceanic (Einarsson 1984).

The substrate in the glacier forefield is characterised by gravel, silt and sand (Jónsson et al. 2016) and the soils are classified as cambic vitrisols and further from the glacier as andosols (Arnalds and Óskarsson 2009).

The area closest to the glacier is mostly unvegetated, with some scattered mosses and lichens. Moss cover (mainly *Racomitrium* sp.) increases with distance from the glacier with 25-50% cover on the oldest moraine (Wojcik et al. 2020).

### 1.1. Sampling

We collected samples of moss and underlying substrate in May 2018. Samples were taken in triplicate along one transect on moraine ridges, at the same locations where Wojcik et al. (2020) collected soil samples (Figure 1).

Moss samples were collected aseptically with a tweezer. Soil samples of 10 cm depth were taken just below the moss cover with a sterilized (with ethanol) hand corer. We collected a total of 27 moss and 27 soil samples. After collection, samples were transported on ice packs for one day and stored at -20 °C until further analyses.

The moss samples were homogenized and split in three parts, one for moss species determination and acetylene reduction assays, one for biogeochemical analysis and one for DNA extraction.

Additional soil samples were taken in late April 2021, to measure pH and moisture. These soil samples were collected at the same coordinates as the samples taken in 2018 and where taken of soil under moss cover and additionally of bare soil. Methods and results of these measurements can be found in Supplementary Methods 1.

### 1.2. Moss shoot length, moisture content and chemical analysis

Moss shoot length was measured for five shoots of each sample. Moss samples were dried at 70°C for 24h and analyzed for field moisture content. Samples were subsequently milled to a fine powder and the total nitrogen (TN) and total carbon (TC) contents and the carbon isotopic composition (δ^13^C) were analyzed. The analysis was carried out at GFZ Potsdam using a mass spectrometer (DELTAplusXL, ThermoFisher) coupled via a ConFlowIII interface with an elemental analyzer (Carlo-Erba NC2500). The analytical precision for δ^13^C was 0.2% and for TC and TN it was 0.01% and replicate determinations showed a standard deviation < 0.02%.

### 1.3. Moss N_2_-fixation rates

Moss N_2_-fixation rates were assessed using the acetylene reduction assay (ARA) method (Hardy et al. 1968). The upper 5 cm of five shoots of each moss sample were weighed and wetted until saturated and then acclimated for 24h at 15 °C in 22 ml vials. Then, we replaced 10% of the headspace (2.2 ml) with acetylene and incubated the samples at 15 °C, under 60 μmol m^−2^ s^−1^ Photosynthetically Active Radiation (PAR) for 24h in a growth chamber (Termaks series 8000, Bergen, Norway). Ethylene and acetylene were quantified by gas chromatography.

Acetylene reduction rates were expressed as ethylene per gram dry weight (field weight) of the moss per day (as in Hardy et al. 1968).

### 1.4. DNA extraction

DNA was extracted for quantification of *nifH* and 16S rRNA gene abundance and 16S rRNA gene sequencing. Before nucleic acid extraction, moss samples were ground in liquid N. DNA from the soil and the moss samples was extracted using the DNeasy PowerSoil Kit (QIAGEN, Hilden, Germany), following the manufacturer’s instructions. DNA concentrations were assessed with a NanoDrop (NanoDrop Technologies, Wilmington, USA).

### 1.5. Quantitative real-time PCR of nifH genes

Quantification of *nifH* genes was performed by quantitative PCR (Corbett Rotor-Gene) using the primer set PolF/PolR. We confirmed the specificity of the *nifH* primers for our samples by Sanger sequencing of 10 clone fragments. Standards for *nifH* reactions were obtained by amplifying one cloned *nifH* sequence with flanking regions of the plasmid vector using the M13 primer sites on the plasmid (TOPO TA cloning Kit, Invitrogen). Standard curves were obtained by serial dilutions (10^6^ to 10^1^ copies per reaction; E = 0.9 – 1.1, R2 = > 0.99 for all reactions).

Each reaction had a volume of 20 µL, containing 10 µL of 2x QuantiFast SYBR Green PCR Master Mix (QIAGEN), 0.2 µL of each primer (10 µM), 0.8 µL of BSA (5 µg/µL), 6.8 µL of RNase free water and 2 µL of template. The cycling program was 5 min at 95 °C, 30 cycles of 10 s at 95 °C and 30 s at 60 °C. Samples with less than 10 *nifH* gene copies per µL and less than 100 16S rRNA gene copies per µL reaction were considered negative.

### 1.6. Sequencing and bioinformatics

Library preparation and paired-end (2 × 300 nt) sequencing of the V3-V4 region of the 16S rRNA gene on an Illumina HiSeq 2500 platform was performed by the Beijing Genomics Institute, using 338F/806R primer pair (Klindworth et al. 2013) and the standard Illumina protocol. We processed the raw sequences using the DADA2 pipeline (Callahan et al. 2016; Callahan, McMurdie, and Holmes 2017), which does not cluster sequences into operational taxonomic units (OTUs), but uses exact sequences or amplicon sequence variants (ASVs). Forward reads were truncated at 250 bp and reverse reads at 220 bp. Assembled ASVs were assigned taxonomy to the SILVA_132 database (Quast et al. 2013) using the Ribosomal Database Project (RDP) naïve Bayesian classifier (Wang et al. 2007) in DADA2. We removed ASVs assigned to chloroplasts and mitochondria and singletons. In total, for 47 samples, 2 972 ASVs remained. To account for uneven sequencing depths, the data were normalized using cumulative-sum scaling (CSS) (Paulson et al. 2013).

### 1.7. Statistics

We used linear models (*lm* from the R package ‘stats’) to investigate the responses of TC, TN, C/N ratio, and moss tissue δ^13^C, N_2_-fixation rates, *nifH* gene abundance, and richness and diversity of the soil and moss associated bacterial communities to moss species and time since deglaciation. We used a post-hoc Tukey test to analyze differences in richness and diversity of bacterial communities in moss species and underlying soil.

To test the effect of time since deglaciation and moss traits on the bacterial community composition of the mosses and the underlying soil, we used PERMANOVAs on weighted Unifrac distance matrices (*adonis* from the R package ‘vegan’). We used moss species as strata in the PERMANOVAs to check whether time since deglaciation and moss traits could explain variation in the bacterial communities in the whole dataset, but we also ran PERMANOVAs on the two moss species separately. To avoid multicollinearity in the linear regression, we only included explanatory factors in the PERMANOVAs with correlation coefficients lower than 0.7 (Table S1).

To identify soil- and moss-associated bacterial taxa whose relative abundance changes with time since deglaciation, we used the R-package ‘DESeq2’ (Love, Huber, and Anders 2014). We used the non-normalised data and an adjusted P-value cut-off of 0.1. For the moss-associated taxa, we included moss species and time since deglaciation in the model (species + time since deglaciation), to correct for moss species, similarly to the linear models.

To explore the direct and indirect relationships between time since deglaciation, moisture content, TN, moss-associated bacterial composition and N_2_-fixation, we constructed a structural equation model (SEM) (using the R package ‘lavaan’ (Rosseel 2012)). For moss-associated bacterial community composition we used the position on the first PCoA axis. As controlling for moss species was not possible, we only used the data from *R. lanuginosum* for the SEM.

## 2. Results

### 2.1. Moss functional traits and N_2_-fixation in the glacier forefield

Moss shoot length increased with time since deglaciation, from 16.5 to 46.9 mm (*P* = 0.02) (Figure 2A, Table S2 and S3). Moss moisture content, TC, TN, C/N ratio and δ13C did not change significantly (Figure 2B-E, Table S2, S4-8), Nevertheless, an non-significant increasing trend with time since deglaciation was found for TC and C/N ratio (Figure 2D and 2E, Table S2, S6 and S7). *nifH* gene abundance decreased (*P* = 0.01) with time since deglaciation (Figure 2G, Table S2 and S9).

**Figure 2.**
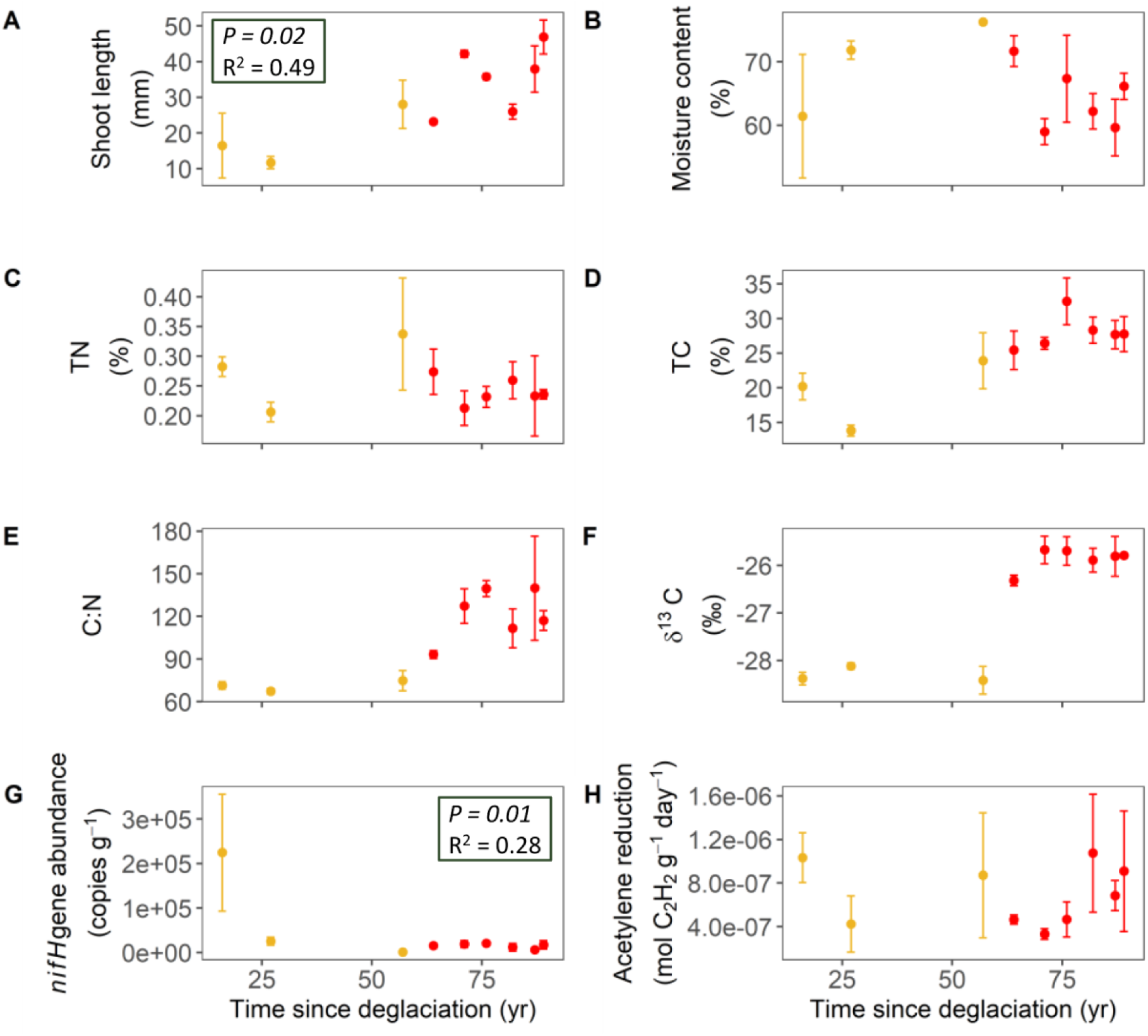
Variations in A) moss shoot length, B) moss moisture content, C) TC, D) TN, E) C/N ratio and F) δ^13^C content of moss shoots, G) *nifH* gene abundance in the mosses and H) moss-associated N_2_-fixation rates (measured by acetylene reduction) with time since deglaciation, in two moss species: *R. ericoides* (yellow) and *R. lanuginosum* (red). Shown are mean ± standard error of triplicate samples from each sampling location. *P*-values and R^2^ are shown in the top left corner of each plot when *P* < 0.05.

Moss δ^13^C was significantly higher in *R. lanuginosum* than in *R. ericoides* (*P* < 0.001) (Figure 2f, Table S8). Moss-associated N_2_-fixation rate (expressed as acetylene reduction rate) showed considerable variation along the chronosequence, but no significant trend with time since deglaciation (Figure 2H, Table S2 and S10). The average acetylene reduction rate in the forefield was 0.00769 µmol C_2_H_2_ Kg^-1^ day^-1^.

### 2.2. Bacterial communities

#### 2.2.1. Richness and diversity of moss and soil bacterial communities

There was no difference in soil or moss microbial diversity between the farthest and oldest edge of our sampling scheme and the closest and more recent land exposed by the glacier retreat (Figure S1, Table S11, S13 and S15). All of the diversity indicators were higher for the soil compared to the two moss species (*P* < 0.001 for both phylogenetic diversity, richness and Shannon diversity; Figure S1, Table S12, S14 and S16).

#### 2.2.2. Moss and soil bacterial community structure

Moss and soil bacterial community composition differed from each other (Permanova R^2^ = 0.41, *P* < 0.001; Figure 3A, Table S17). The soil bacterial community changed significantly with time since deglaciation (Permanova R^2^ = 0.10, *P* = 0.04; Figure 3C, Table S18). Part of the variation in the structure of the soil bacterial communities was related to moss C/N ratio (Permanova R^2^ = 0.09. *P* = 0.04; Figure 3C, Table S18). The moss bacterial community (regardless of moss species) was also affected by time since deglaciation (Permanova R^2^ = 0.38, *P* < 0.001) and additionally by moisture content (Permanova R^2^ = 0.12, *P* < 0.001; Table S19).

**Figure 3.**
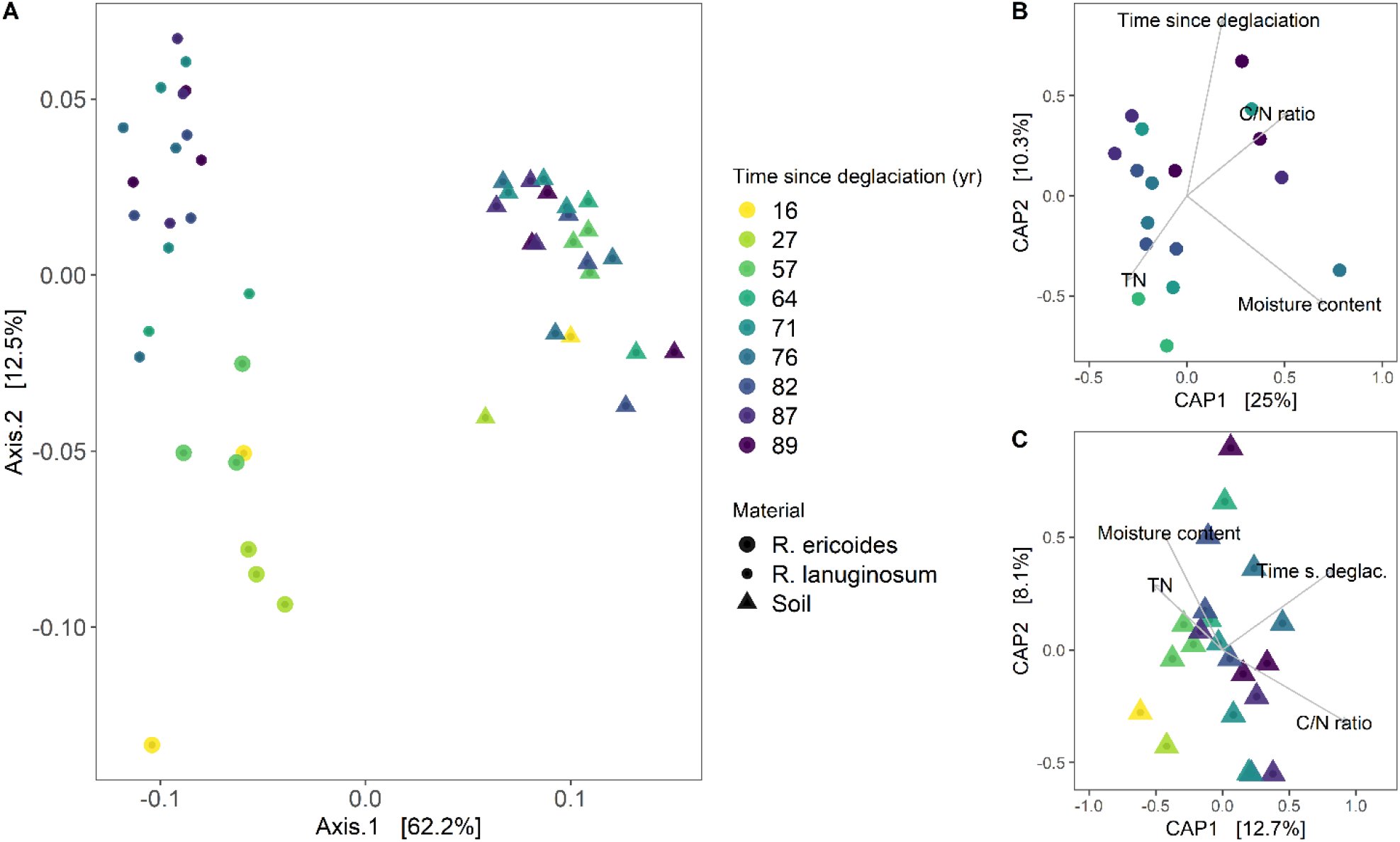
A) Principal coordinate analysis (PCoA) biplot of the bacterial communities of the mosses *R. ericoides* and *R. lanuginosum* and the underlying soil on ASV level based on weighted unifrac distances and B) CAP analysis of the bacterial community of the moss *R. lanuginosum* and environmental factors and C) CAP analysis of the underlying soil bacterial community environmental factors.

When moss species were analyzed separately, the Permanova showed that the *R. ericoides* bacterial community changed with time since deglaciation (Permanova R^2^ = 0.22, *P* = 0.004), but was also affected by moss C/N ratio (Permanova R^2^ = 0.18, *P* = 0.02) and moss moisture content (Permanova R^2^ = 0.24, *P* = 0.003; Table S20). The *R. lanuginosum* bacterial community was not affected by time since deglaciation, but varied with moss moisture content (Permanova R^2^ = 0.19, *P* = 0.003; Figure 3B, Table S21).

#### 2.2.3. Bacterial community composition

The bacterial communities of the mosses in the Fláajökull glacier forefield were characterised by Proteobacteria (35% and 28% on average in *R. ericoides* and *R. lanuginosum* respectively), Acidobacteria (15% and 23%), Bacteroidetes (25% and 24%), Verrucomicrobia (7% and 7%), Chloroflexi (1% and 5%) and Actinobacteria (10% and 7%), on phylum level (Figure 4). Cyanobacteria abundance was relatively low in the moss species (3% and 2%).

**Figure 4.**
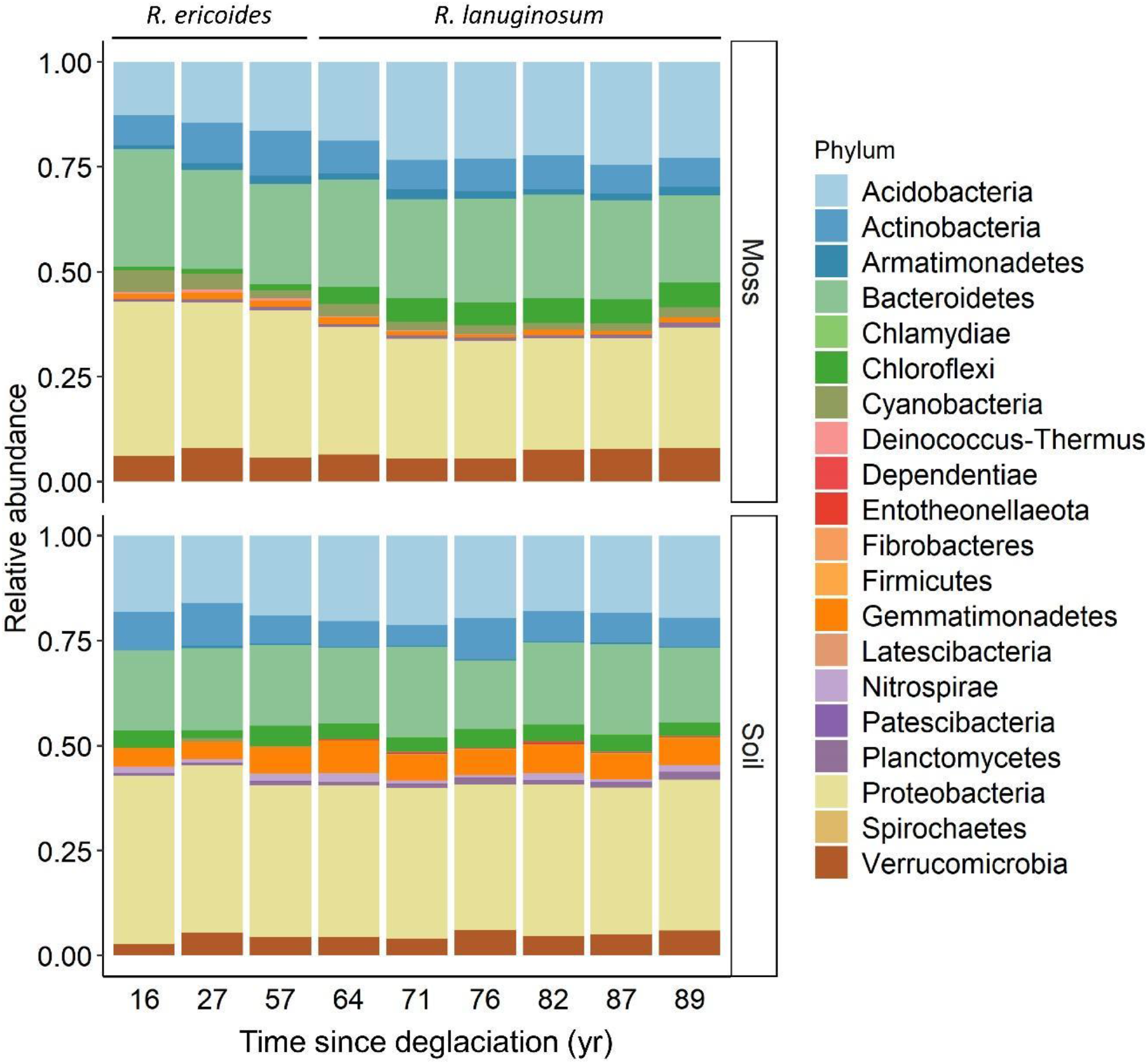
Phyla-level composition of the bacterial communities of *Racomitrium* mosses and underlying soil along a chronosequence in the Fláajökull glacier forefield.

Alphaproteobacteria were the most abundant class within the Proteobacteria (22% and 19%). The Alphaproteobacteria were dominated by the families Acetobacteraceae (7% and 8%) and Sphingomonadaceae (7% and 4%) (Figure S2). Acidobacteria were dominated by the families Acidobacteriaceae (6% and 13%) and Solibacteraceae (6% and 8%) (Figure S3). There were no clear dominant families within the Actinobacteria (Figure S3). Within this phylum, we identified Frankiaceae (1% in both *R. ericoides* and *R. lanuginosum*), Ilumatobacteraceae (1% in both *R. ericoides* and *R. lanuginosum*), Nakamurellaceae (2% and <1%) and Pseudonocardiaceae (1% in both *R. ericoides* and *R. lanuginosum*) in all successional stages (Figure S4). Bacteroidetes were dominated by the family Chitinophagaceae (14% and 15%) (Figure S5) and Cyanobacteria were dominated by the genus *Nostoc* (3% and 2%) (Figure S6).

The soil bacterial communities were dominated by the phyla Proteobacteria (36%), Acidobacteria (19%), Actinobacteria (7%) and Bacteroidetes (19%) (Figure 4).

The classes Alpha- and Gammaproteobacteria showed similar abundances in all soil samples along the chronosequence (15% and 16% for Alpha- and Gammaproteobacteria respectively). The Alphaproteobacteria were dominated by Xanthobacteraceae (5%) and Sphingomonadaceae (2%) (Figure S2). The family Nitrosomonadaceae (5%) dominated the Gammaproteobacteria (Figure S7). The Acidobacteria were dominated by the families Solibacteraceae (7%), Pyrinomonadaceae (7%) and Blastocatellaceae (4%) (Figure S3). The abundance of Actinobacteria was similar to their abundance in the moss samples, relatively variable and lacking a clearly dominating taxon. The Ilumatobacteraceae (1%) occurred in all soil samples (Figure S4). The Bacteroidetes were dominated the family Chitinophagaceae (11%) (Figure S5).

#### 2.2.4. Bacterial community composition across time since deglaciation and moss species

On phylum level, the relative abundance of Chloroflexi increased across our chronosequence, both in moss and soil; while Proteobacteria, Cyanobacteria and Bacteroidetes decreased in the moss (Figure 4, Figure S8 and S9).

On ASV level, we detected more ASVs changing in relative abundance with time since deglaciation in the soil than in the moss (Figure 5). All detected ASVs increased in relative abundance across the chronosequence. Most of these ASVs belonged to the Proteobacteria. The two ASVs that increased with time since deglaciation belonged to the candidate genus Solibacter and the family Acetobacteraceae. The ASVs showing the strongest increase in relative abundance with time since deglaciation in the soil belonged to the families Acetobacteraceae, Micropepsaceae and Chinitophagaceae and the genera *Parafilimonas* and *Nocardioides*.

**Figure 5.**
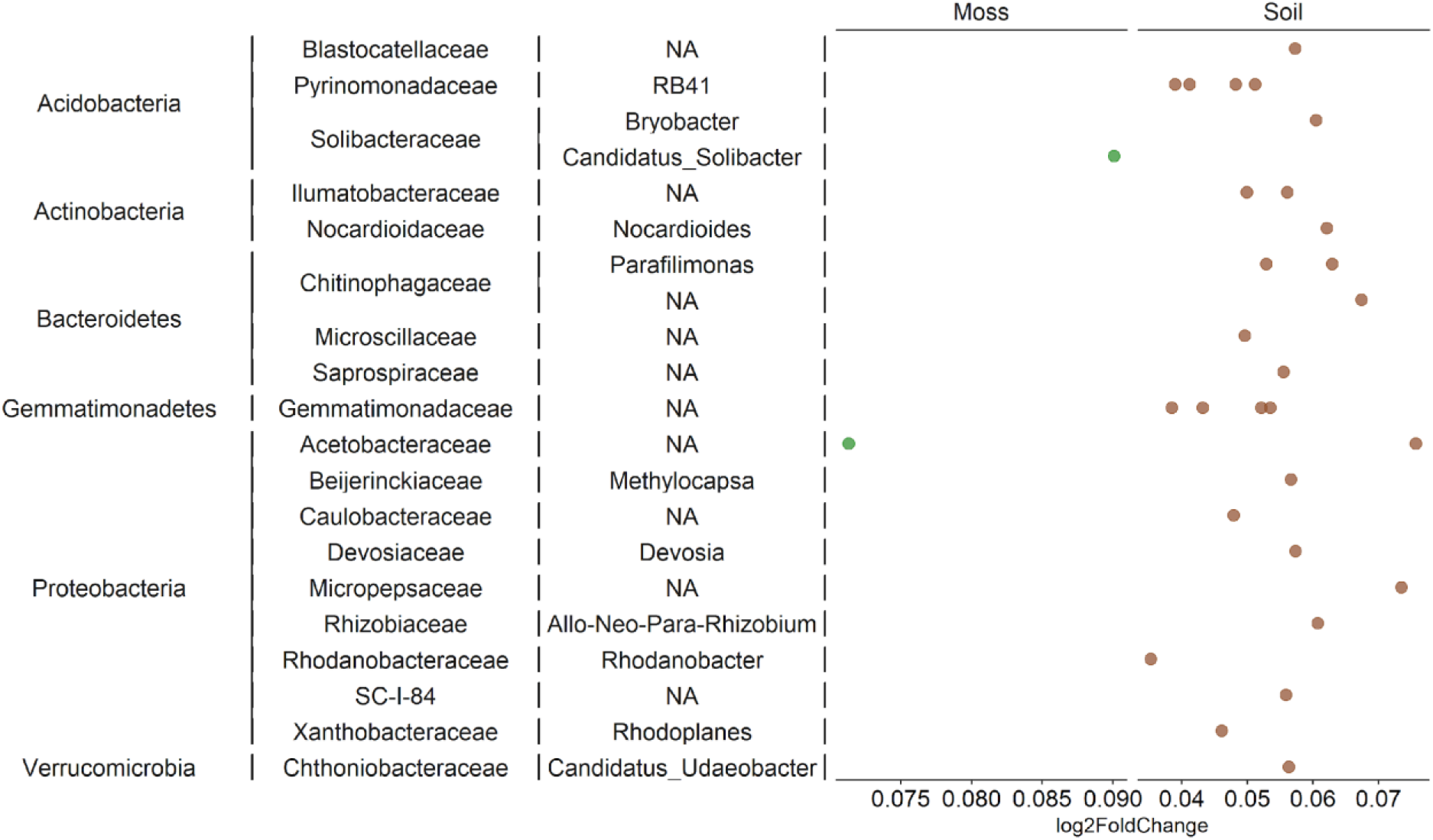
Changes in relative abundance of microbial groups (at ASV level) across the chronosequence in *Racomitrium* moss species (green) and underlying soil (brown).

#### 2.2.5. Linkages between N_2_-fixation, time since deglaciation, moisture content, TN and bacterial community structure

We used structural equation modeling to investigate the direct and indirect linkages between time since deglaciation, moisture content, moss TN and TC, the *R. lanuginosum*-associated bacterial community structure, *nifH* gene abundance and N_2_-fixation (Figure 6 and Table S22). Bacterial community structure was positively affected by TN (standardized path coefficient 0.41 *P* < 0.01) and negatively affected by moisture content (standardized path coefficient -0.63; *P* < 0.001)). *nifH* gene abundance was negatively affected by the structure of the bacterial community (standardized path coefficient -0.87; *P* < 0.001) and N_2_-fixation rate was negatively affected by *nifH* gene abundance (standardized path coefficient -0.68; *P* = 0.01). This indirect effect of the bacterial community structure on N_2_-fixation rates via changes in *nifH* gene abundance was also significant (standardized path coefficient 0.60; *P* = 0.047; Table S22).

**Figure 6.**
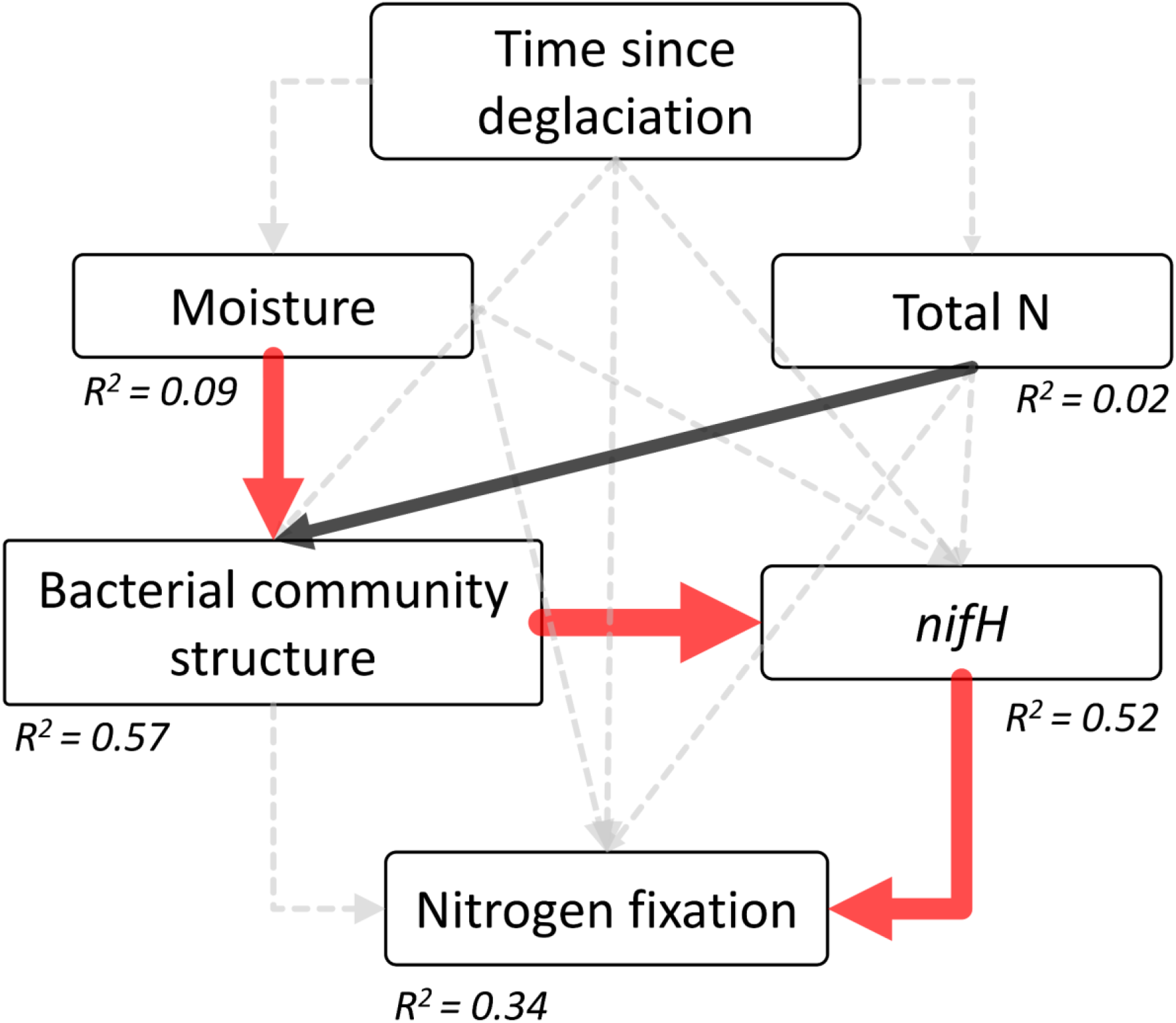
Structural equation model (SEM) showing linkages between time since deglaciation, moss moisture content, moss total N and moss bacterial community and N_2_-fixation. Note that this SEM is only based on the moss *R. lanuginosum*. χ^2^ = 2.59, P = 0.459, df = 3.00, GFI = 0.975, RMSEA = 0, TLI = 1.031. Positive significant effects are represented in black and significant negative significant effects in red. Non-significant effects are indicated with dash-line arrows. The strength of the effect is visualized by the width of the arrow. The R^2^-value represents the proportion of total variance explained for the specific dependent variable. Standardized path coefficients are presented in Table S22.

## Discussion

Mosses are among the first colonising plants on newly exposed substrates following glacier retreat. Mosses and their bacterial communities play important roles in the C and N cycle, which are crucial during ecosystem development in glacier forefield. But it is unclear how moss bacterial communities develop during primary succession. Here, we studied moss traits, moss-associated bacterial communities and N_2_-fixation as well as the bacterial communities of the underlying substrate along a chronosequence in the Fláajökull glacier forefield in Iceland. We found links between time since deglaciation and moss traits and bacterial community composition of the mosses and the underlying soil. We also found that soil and moss bacterial community composition are related to certain moss traits. N_2_-fixation rates were linked to bacterial community structure, but not time since deglaciation. Our new data set on primary succession as a driver of moss-associated bacterial community composition and associated processes contributes to the understanding of biogeochemical cycling in newly exposed ice-free substrates.

### Changes in moss functional traits with time since deglaciation

We hypothesized that moss shoot traits would change time since deglaciation.. Most of the changes occurred in the earlier stages of the successional gradient and stabilised in the later stages. We expected TC to increase with succession, and the overall pattern pointed in that direction albeit not significantly, at least until 76 year after deglaciation. This TC trend in the moss partly agrees with the patterns of soil organic carbon content (SOC) in the same Fláajökull forefield in a parallel study by Wojcik et al. (2020), where the authors showed that TC content increased until the 1936 moraine and that TC was lower on the 1931 and 1929 moraines, probably due to soil disturbance via geomorphological events or due to the heterogeneous nature of the soil substrates in the Fláajòkull forefield.

We also expected moss TN to increase with time since deglaciation, but our data did not show any successional trends in moss N content. In the early successional stages of the Fláajökull forefield soil TN increases (Wojcik et al. 2020), but moss shoot TN didn’t correlate with the soil TN trend. There are several potential reasons for the discrepancy between moss and soil TN. Moss N may for instance be lost via denitrification or via leeching to deeper soil layers (Johnson, Neuer, and Garcia-Pichel 2007). Moss N has also been found to be more rapidly lost from moss litter than C during decomposition (Philben et al. 2018). Nevertheless, as moss mat coverage and shoot length increased with succession, moss TN per m^2^ will increase. Overall moss C/N showed an increasing trend along the chronosequence, probably driven by the increasing C content, but again with lower values on the three oldest, potentially disturbed soils. The increase is similar to the increase in C/N found in the bryophytes with succession in glacier forefields in Tierra del Fuego in Chile (Arróniz-Crespo et al. 2014).

δ^13^C can reflect the signal of multiple environmental factors (Waite and Sack 2011) and often increases in moss tissue with ecosystem age (Bansal, Nilsson, and Wardle 2012; Jonsson et al. 2015). Our data did show less negative values in moss shoot δ^13^C with time since deglaciation. This however could also be due to differences between moss species, with lower values in *R. ericoides* (−28.3 ‰ ± 0.1) and more negative values in *R. lanuginosum* (−25.8 ‰ ± 0.1). This confirms the importance of moss species for δ^13^C values (Bramley-Alves et al. 2015; Waite and Sack 2011). Our average δ^13^C value for *R. lanuginosum* (−25.8 ‰ ± 0.1) is comparable to those found in *R. lanuginosum* on Mauna Loa, Hawaii (−26.3% ± 0.4) (Waite and Sack 2011).

### Potential drivers of the moss-associated and underlying soil bacterial community structure

The bacterial community structure of the mosses and the soil were both affected by time since deglaciation. As moss species an important factor for the composition of the bacterial communities (Holland-Moritz et al. 2018; Bragina et al. 2012), the shift in moss species along the chronosequence may also have contributed to the effect of time since deglaciation on the moss bacterial communities.

Moss moisture content turned out to be an important factor contributing to variation in the bacterial community structure of both *R. ericoides* and *R. lanuginosum*. Moisture content is an important driver of microbial decomposition (Schimel et al. 1999) and may thereby also affect bacterial community structure, especially in the decomposing part of the moss shoots. Interestingly, moisture has also been found to affect the occurrence of Antarctic moss-associated fungi (Hirose et al. 2016). In our study time since deglaciation and C/N ratio affected the bacterial community composition of *R. ericoides*, but not *R. lanuginosum*. The discrepancy in factors structuring the bacterial communities of the two mosses may also be caused by the smaller sample size of *R. ericoides* versus *R. lanuginosum*, but factors driving moss bacterial communities may also change with succession. Our results indicate that time since deglaciation and C/N ratio are important in the earlier stages of succession (eg. in *R. ericoides*), potentially because C/N ratio becomes more stable in the later stages (eg. in *R. lanuginosum*).

The soil bacterial community structure below the mosses showed variation with succession, but less than the overall moss bacterial community. Interestingly, moss C/N ratio was also a driver of the soil bacterial community composition. Moss C/N ratio may influence soil C/N ratio, without directly affecting the soil bacterial community, but plant traits such as leaf N can influence soil bacterial community composition (de Vries et al. 2012) and moss chemical traits may thus also affect bacterial community in the underlying substrate.

The bacterial communities of *Racomitrium* moss species and underlying soil were clearly distinct at the ASV level, indicating that there might be little or no lateral transmission of the soil bacterial communities to the moss bacterial communities and/or vice versa. The bacterial community of *R. lanuginosum* in the Fláajökull glacier forefield was also similar to the bacterial community of *R. lanuginosum* from a subarctic-alpine heathland in northwest Iceland (Klarenberg et al. 2021). Many taxa are shared in similar proportions, such as the orders Acetobacterales, Acidobacterales and Solibacterales, while Bacteroidetes were more abundant in the mosses in the glacier forefield and Planctomycetes more abundant in the heathland. Cyanobacteria were less abundant in the moss in the glacier forefield than in the heathland, potentially because the Fláajökull glacier forefield receives more N from deposition as the forefield is in the vicinity of farmlands that could provide the moss with N and reduce the need for Cyanobacteria as diazotrophic symbionts. Generally, the moss-associated bacterial communities are dominated by presumptively acidophilic bacteria often associated with ombrotrophic or other oligotrophic environments and are comparable to the bacterial communities of other moss species (Holland-Moritz et al. 2021).

### Taxa specific trajectories in of moss-associated and underlying soil bacterial communities with succession

Most of the bacterial phyla that shifted in relative abundance during succession were found in the moss. The phylum Chloroflexi increased in relative abundance with succession in both the soil and the mosses. An increase in Chloroflexi with succession has also been detected in the soil and rhizosphere of *Saxifraga oppositifolia* in a glacier forefield in the high Arctic (Mapelli et al. 2018). In the moss, we found decreasing abundance of Proteobacteria, Cyanobacteria and Bacteroidetes. These taxa often become less abundant as succession progresses in glacier forefields in soils (Bajerski and Wagner 2013; Jiang et al. 2018; Fernández-Martínez et al. 2017; Bradley et al. 2016) and our results show that similar patterns are found in the moss microbiome, but less so in the moss-covered soil.

On ASV level, most changes with soil age were detected in the soil bacterial community. All of these ASVs increased in relative abundance with soil age. Many of them were classified as genera known to be able to degrade plant-organic matter, such as *Ca. Solibacter* (Ward et al. 2009), *Nocardioides* (Guo et al. 2021), members of the Chitinophagaceae (Yong Li et al. 2011) and Micropepsaceae (Harbison et al. 2016), indicating increased moss abundance with succession also increases the potential for degradation of dead moss material. While Cyanobacteria decreased in relative abundance with soil age in the moss, heterotrophic N_2_-fixers became more abundant with soil age (for instance *Devosia* (Rivas et al. 2002), Rhizobiaceae (Dobbelaere, Vanderleyden, and Okon 2003), Methylocapsa (Dedysh et al. 2002) and *Rhodoplanes* (Buckley et al. 2007) in soil, and Acetobacteriaceae (Saravanan et al. 2008) in the mosses), probably because of increased substrate availability. An increase in potential denitrifiers (*Ca. Solibacter* (Ward et al. 2009) and *Rhodanobacter* (Kostka et al. 2012)) suggests an increase in nitrates and/or nitrites with succession and loss of N via denitrification with succession. Some of the taxa increasing along the chronosequence are known to be acidophilic (Chitinophagaceae and Gemmatimonadaceae (Cline and Zak 2015), Acetobacteraceae in moss (Kersters et al. 2006), Micropepsaceae (Harbison et al. 2016)), and may be linked to decreased soil pH with soil age in the Fláajökull glacier forefield ((Wojcik et al. 2020) and Table S24).

### Moss N_2_-fixation and diazotroph abundance during succession

We did not detect any trends in N_2_-fixation rates with soil age. *nifH* gene abundance however, showed an overall decrease with soil age, indicating a decreasing abundance of diazotrophs with succession. Moss-associated N_2_-fixation rates were not affected by moss N content, soil age or moisture, but rather by the abundance of diazotrophs and bacterial community composition, at least in *R. lanuginosum*. The negative link between *nifH* gene abundance and N_2_-fixation rates, could indicate that not all bacteria taxa possessing *nifH* genes are actively involved in N_2_-fixation, or that the degeneracy of the *nifH* gene primer pair PolF/PolR is not high enough to target most N_2_-fixing taxa in the moss bacterial communities. N_2_-fixation rates could also depend on bacterial community composition. The last explanation is supported by the indirect connection found between bacterial community composition and N_2_-fixation rates via *nifH* gene abundance. We for instance detected a decrease in the relative abundance of Cyanobacteria in the mosses with soil age and an increase in an ASV of the Acetobacteraceae, which contain nitrogen fixing members (Saravanan et al. 2008). Additionally, past research has shown that shifts in *nifH* gene diversity with succession occur in soil in glacier forefields (Duc et al. 2009) and our study suggests that these shifts may also take place in mosses.

Moss N_2_-fixation may be and stay an important source of N throughout glacier forefields, with increasing importance as moss cover increases with succession. The relative importance of N_2_-fixation and mineralization for the N content of the moss and the soil along the chronosequence may be better understood when ^15^N depletion is taken into account in future studies.

## 3. Conclusion

We studied the development of *Racomitrium* moss bacterial communities as well as those of the underlying substrate in relation to moss functional traits along a chronosequence in the glacier forefield of Fláajökull in southeast Iceland. While moss functional traits such as TN and moisture content did not show clear trends along the chronosequence, moss shoot length increased with succession. Time since deglaciation as well as moss C/N ratio and moss moisture content were related to moss bacterial community structure, showing for the first time how moss functional traits are important drivers for moss-associated bacterial communities. The bacterial communities of the underlying soil were also affected by time since deglaciation and by moss C/N ratio, highlighting the influence of moss cover on soil development. Moss and underlying soil bacterial communities differed strongly from each other, suggesting that little lateral transfer between them takes place. We did not detect any trends in moss-associated N_2_-fixation rates with time since deglaciation or moss TN, but N_2_-fixation rates were linked to bacterial community structure and negatively linked to *nifH* gene abundance. This may indicate a shift in diazotrophic taxa with different N_2_-fixing efficiencies along the chronosequence and our data indeed show a proportional decrease in Cyanobacteria and an increase in heterotrophic N_2_-fixing taxa.

Our study underlines the importance of moss functional traits as potential drivers for moss bacterial community structure, but also links moss functional traits to bacterial communities in the underlying substrate. This is one way in which mosses can enhance soil development in glacier forefields, but our results also show that moss-associated N_2_-fixation takes place along the whole chronosequence and thereby likely contributes to N availability. Our study contributes to the understanding of the role of mosses in ecosystem development, which will be increasingly important in a future warmer climate leading to increased glacier retreat.

## Acknowledgements and funding

This work was supported by the MicroArctic Innovative Training Network grant supported by the European Commission’s Horizon 2020 Marie Sklodowska-Curie Actions program [grant number 675546] and the University of Akureyri Science Fund [grant number R1910].

We kindly thank Dr. Birgit Plessen for the TC, TN and δ^13^C measurements. We would also like to thank the Tundra Ecology group at the University of Iceland for interesting discussions and helpful advice for the data analysis.

## Author contributions

IJK designed the study, OV and IJK collected the samples. IJK, CK and AS performed the laboratory analysis. IJK analysed the data and wrote the paper with input from OV, CK, AS and LGB.

## Data availability

Raw sequences are available in the European Nucleotide Archive under accession number XXXX.

