## Supplementary Material for "Moss functional traits are important drivers for moss and underlying soil bacterial communities: evidence from a chronosequence in an Icelandic glacier forefield"

**Figure S1.** ASV (Amplicon Sequence Variant) richness, Shannon diversity and Faith's phylogenetic diversity of the bacterial communities of the mosses *R. ericoides* and *R. lanuginosum* and underlying soil along the chronosequence. Shown are mean ± standard error for three replicates from each moss species and soil collected at each sampling point.

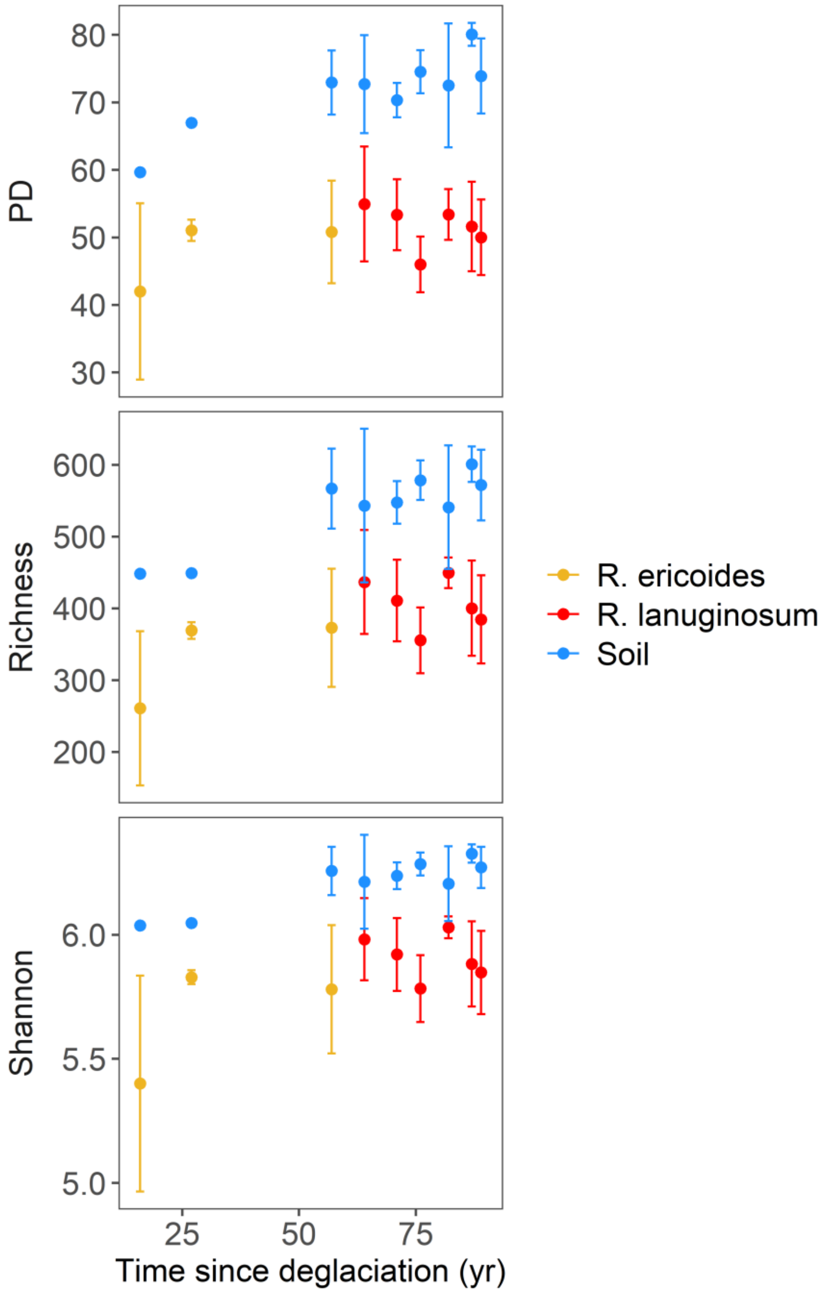

**Figure S2.** Family level composition of the Alphaproteobacteria within the bacterial communities of *Racomitrium* mosses and underlying soil along a chronosequence in the Fláajökull glacier forefield.

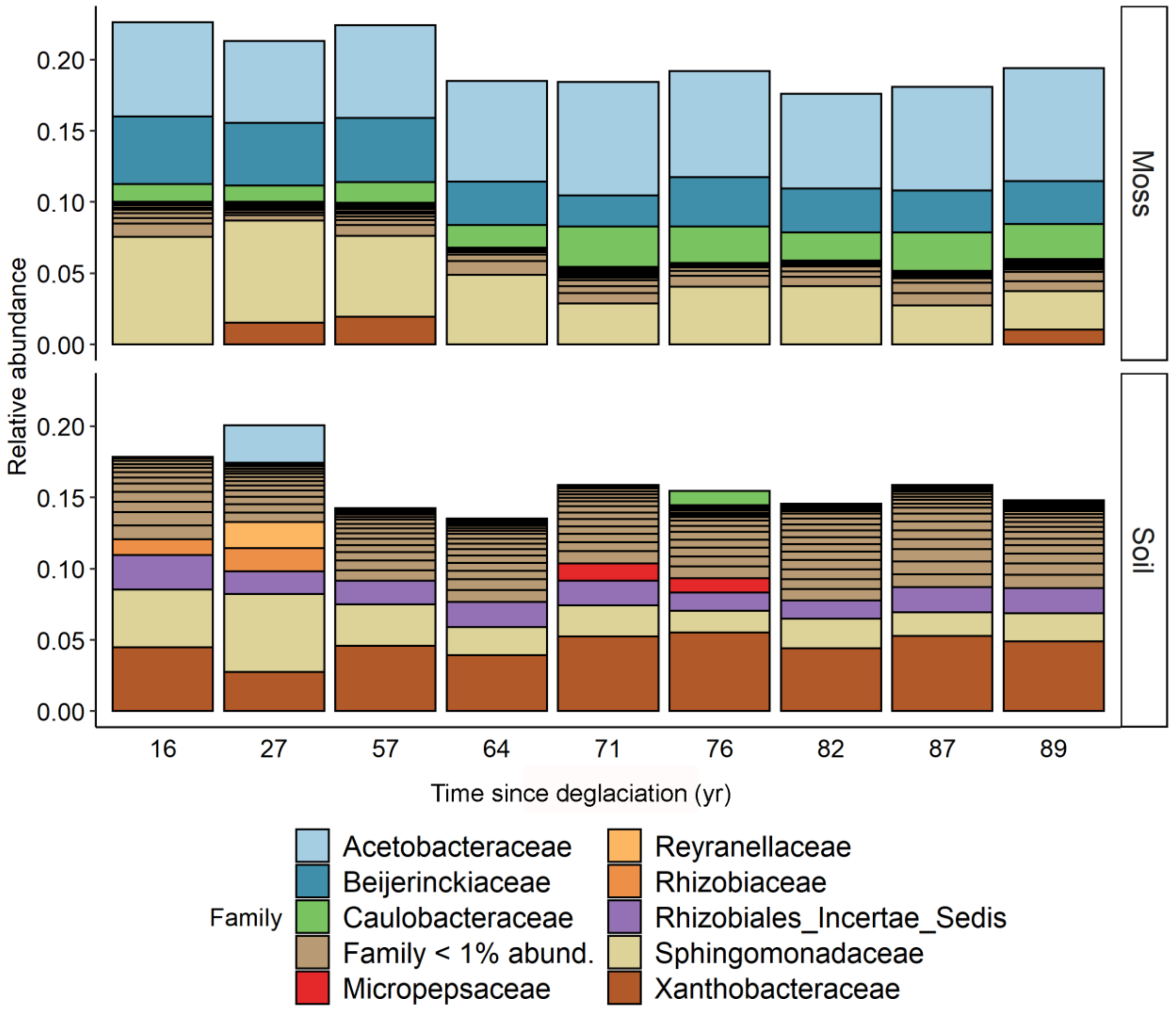

**Figure S3.** Family level composition of the Acidobacteria within the bacterial communities of *Racomitrium* mosses and underlying soil along a chronosequence in the Fláajökull glacier forefield.

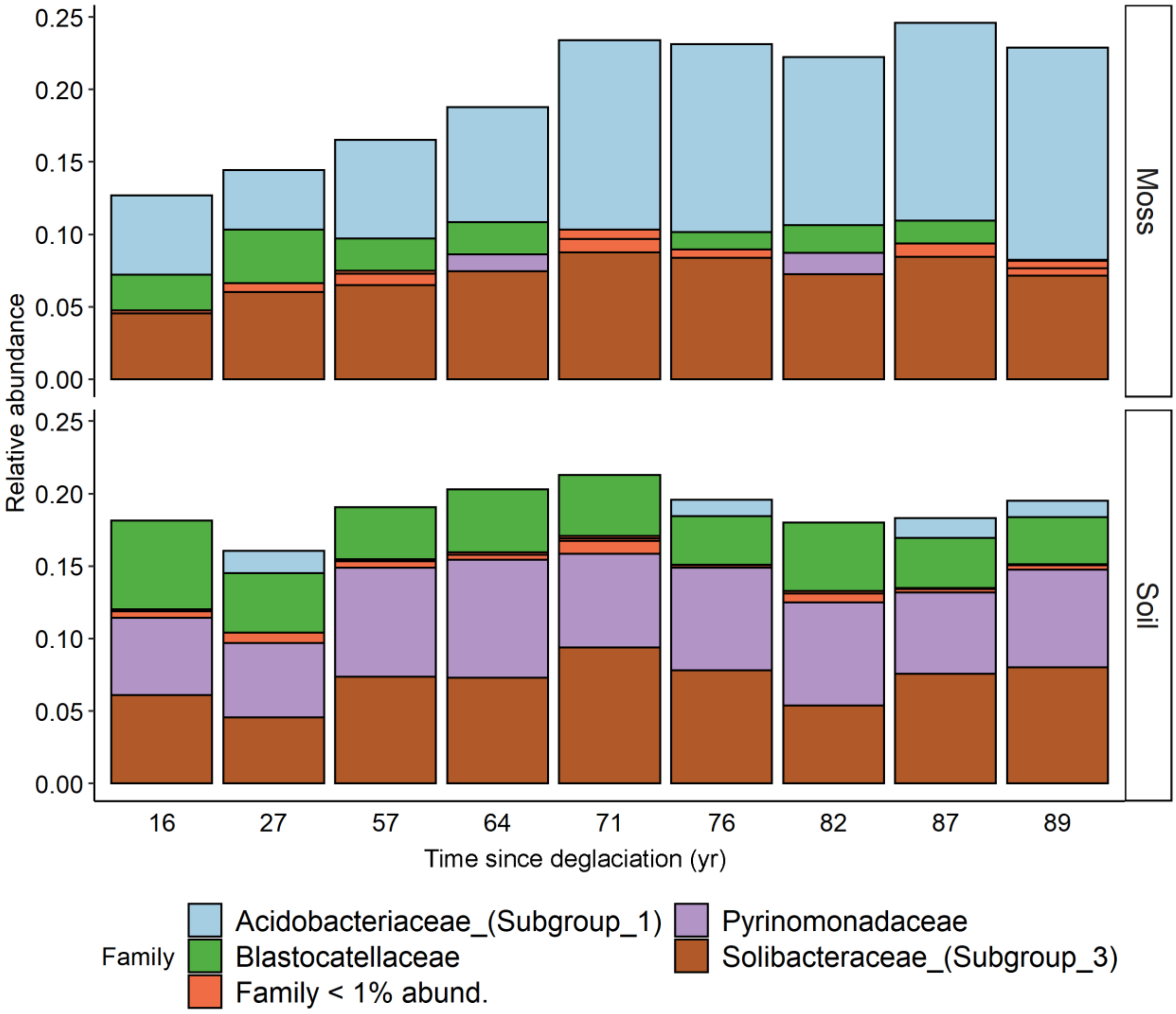

**Figure S4.** Family level composition of the Actinobacteria within the bacterial communities of *Racomitrium* mosses and underlying soil along a chronosequence in the Fláajökull glacier forefield.

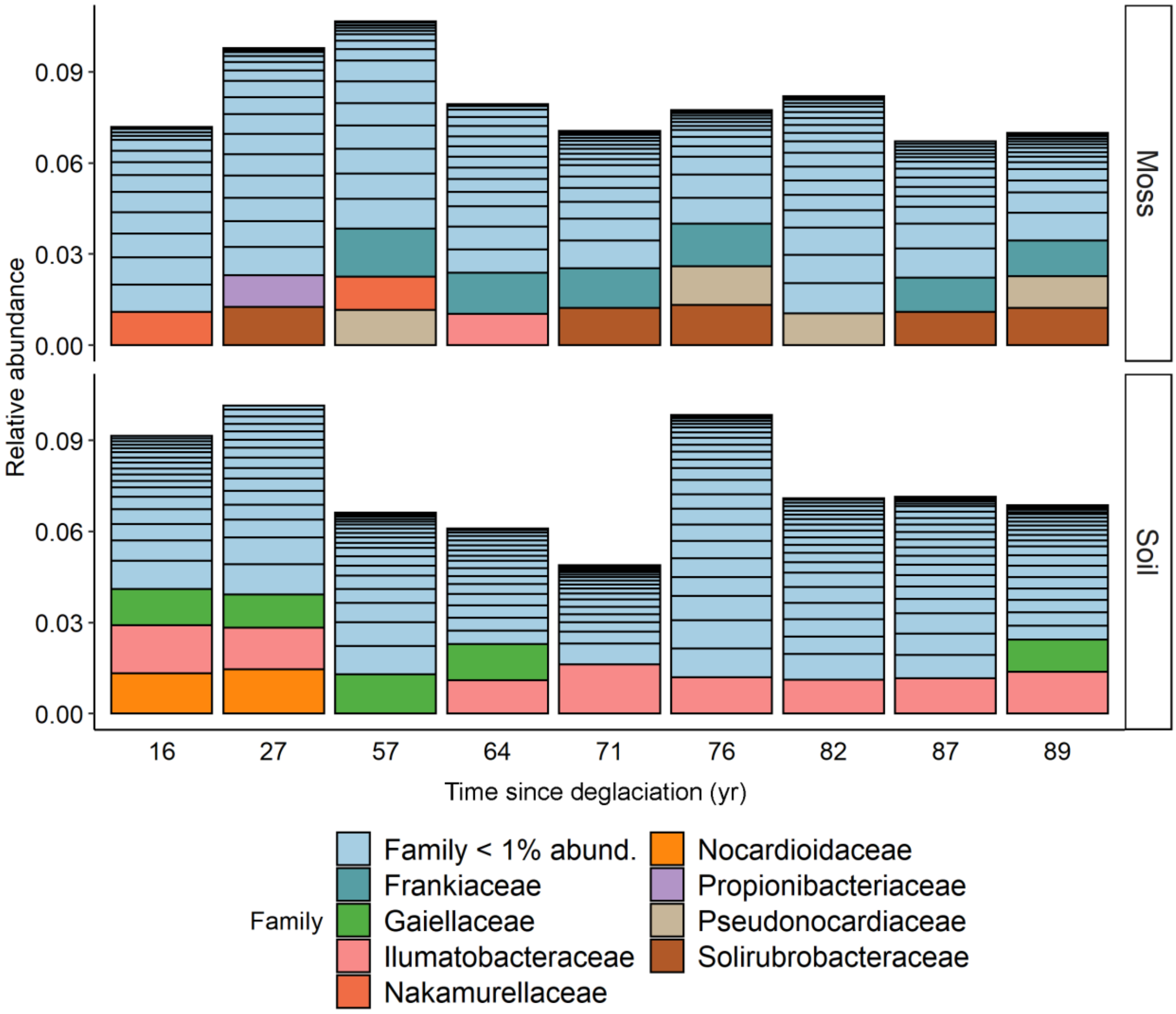

**Figure S5.** Family level composition of the Bacteroidetes within the bacterial communities of *Racomitrium* mosses and underlying soil along a chronosequence in the Fláajökull glacier forefield.

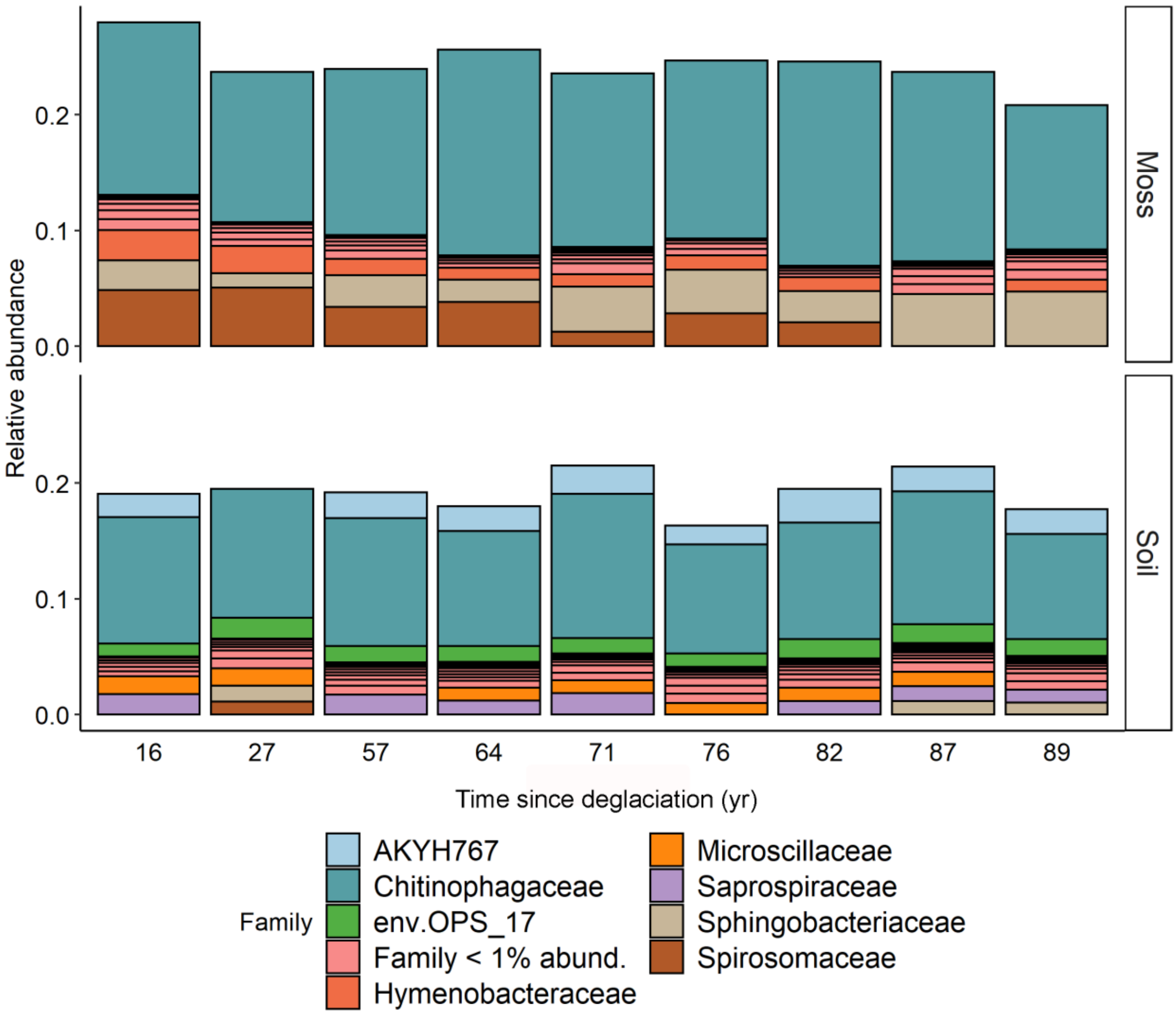

**Figure S6.** Genus level composition of the Cyanobacteria within the bacterial communities of *Racomitrium* mosses and underlying soil along a chronosequence in the Fláajökull glacier forefield.

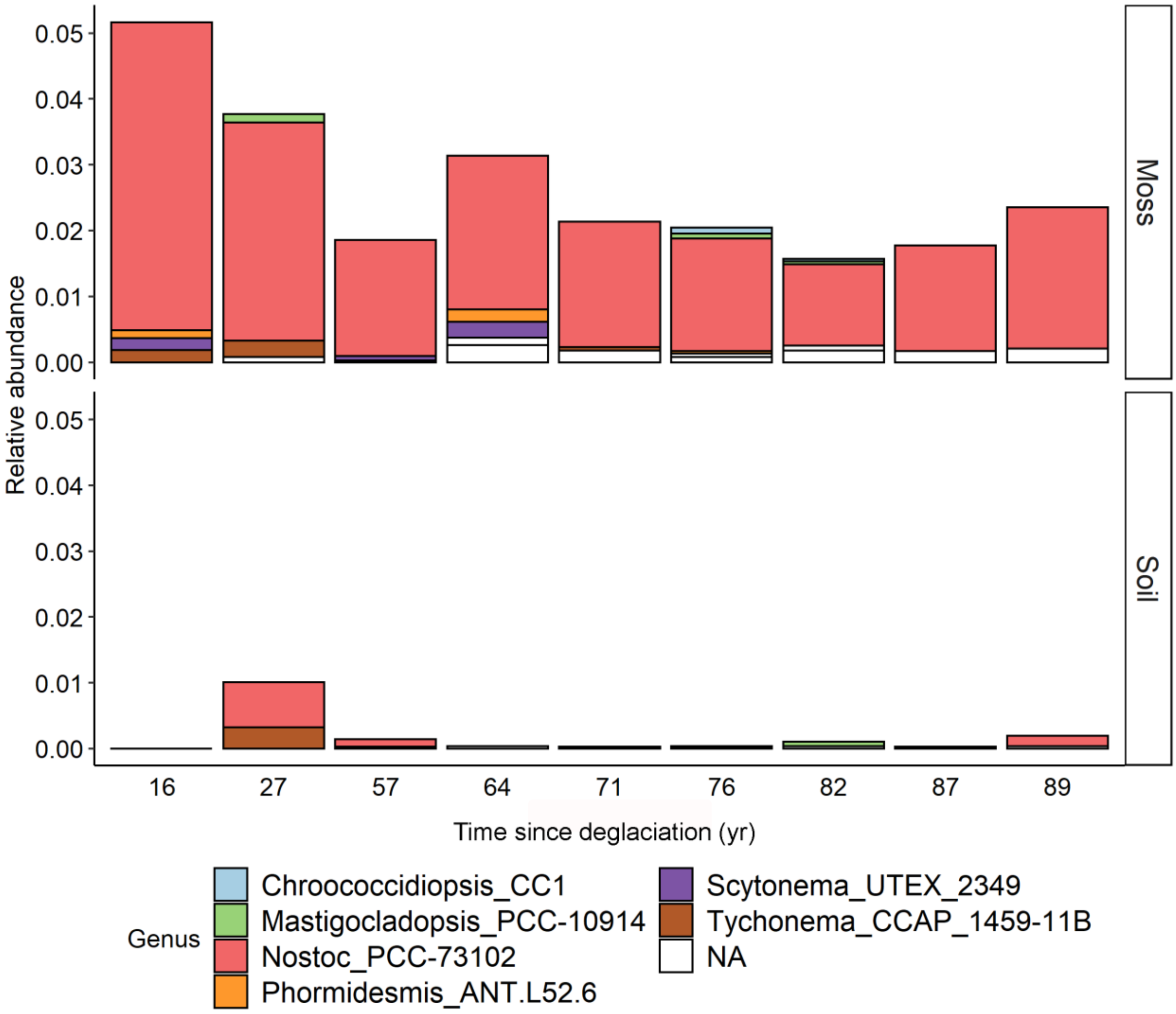

**Figure S7.** Family level composition of the Gammaproteobacteria within the bacterial communities of *Racomitrium* mosses and underlying soil along a chronosequence in the Fláajökull glacier forefield.

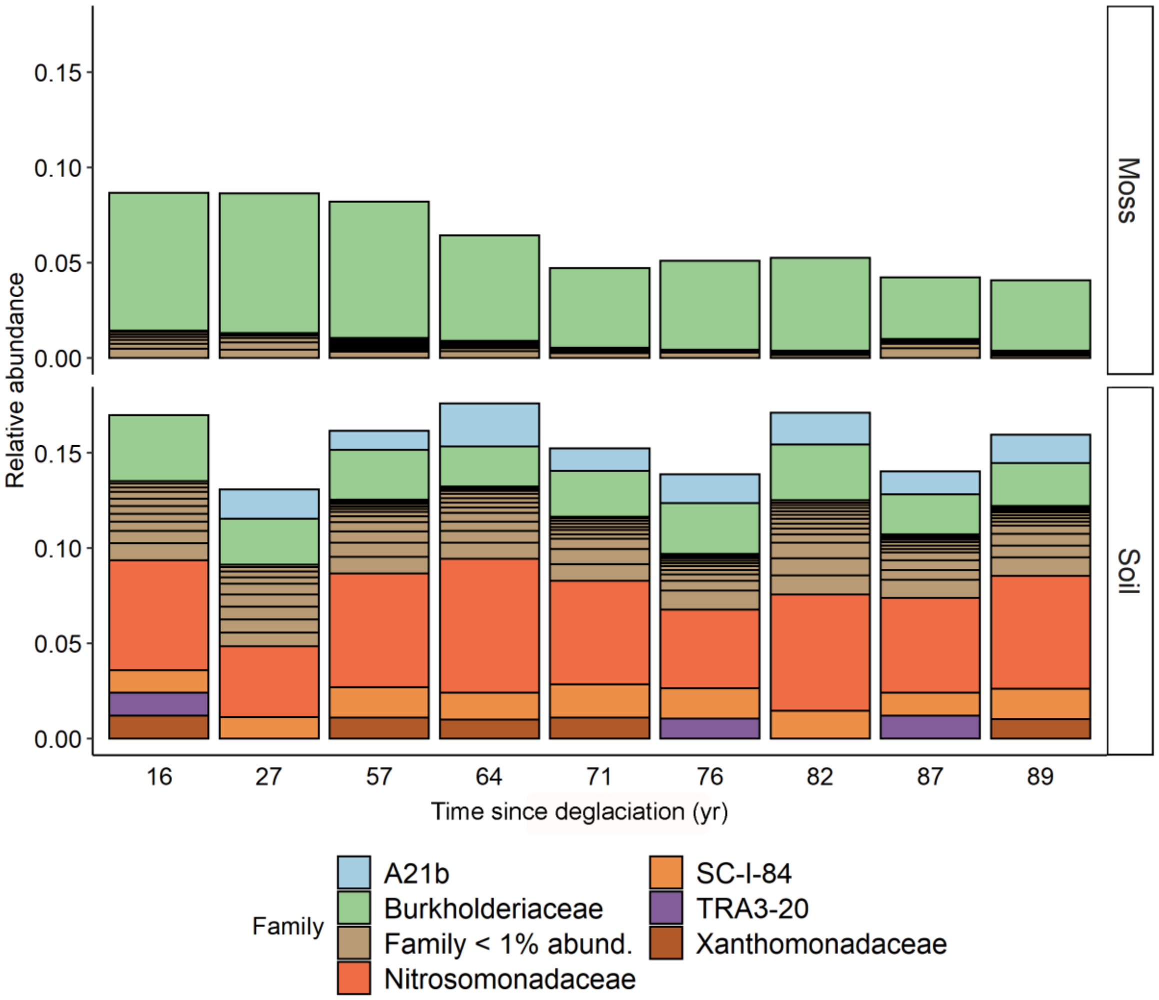

**Figure S8.** Bacterial phyla changing in relative abundance with soil age in the moss detected by DESeq2.

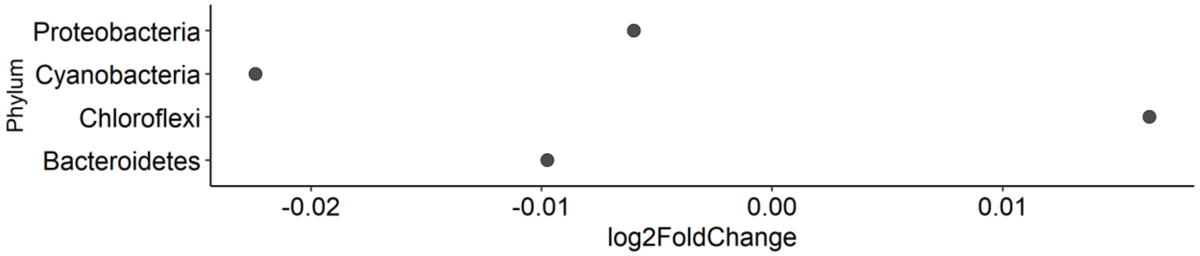


**Figure S9.** Bacterial classes changing in relative abundance with soil age in the soil detected by DESeq2.

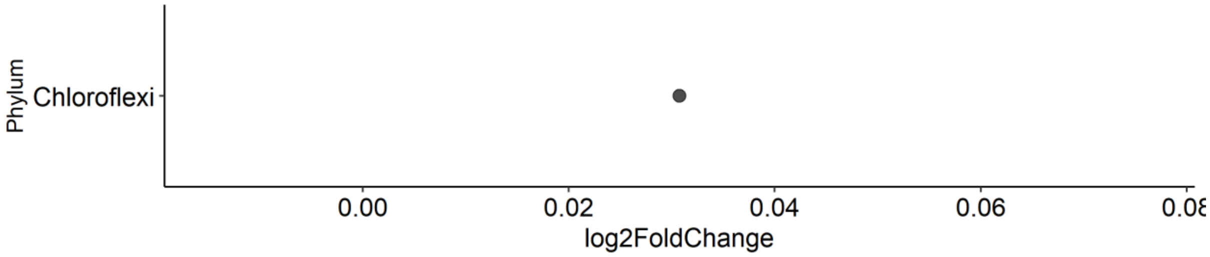


**Figure S10** Soil moisture content (%) of bare soil and moss-covered soil along the chronosequence in the Fláajökull glacier forefield.

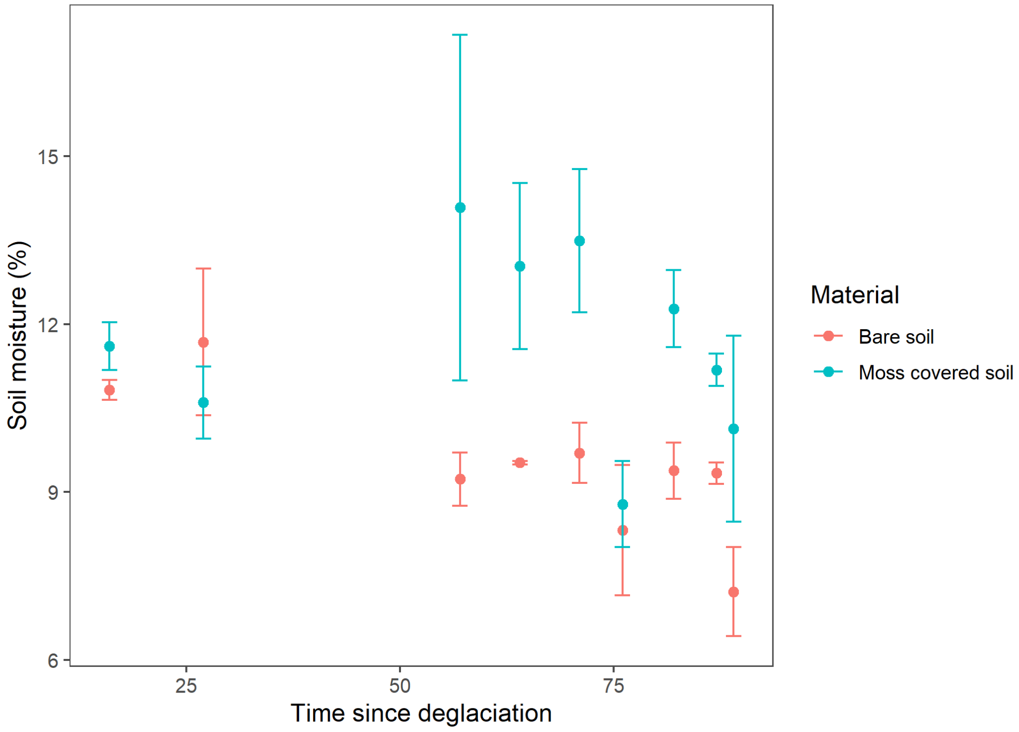

**Figure S11** pH of bare soil and moss-covered soil along the chronosequence in the Fláajökull glacier forefield.

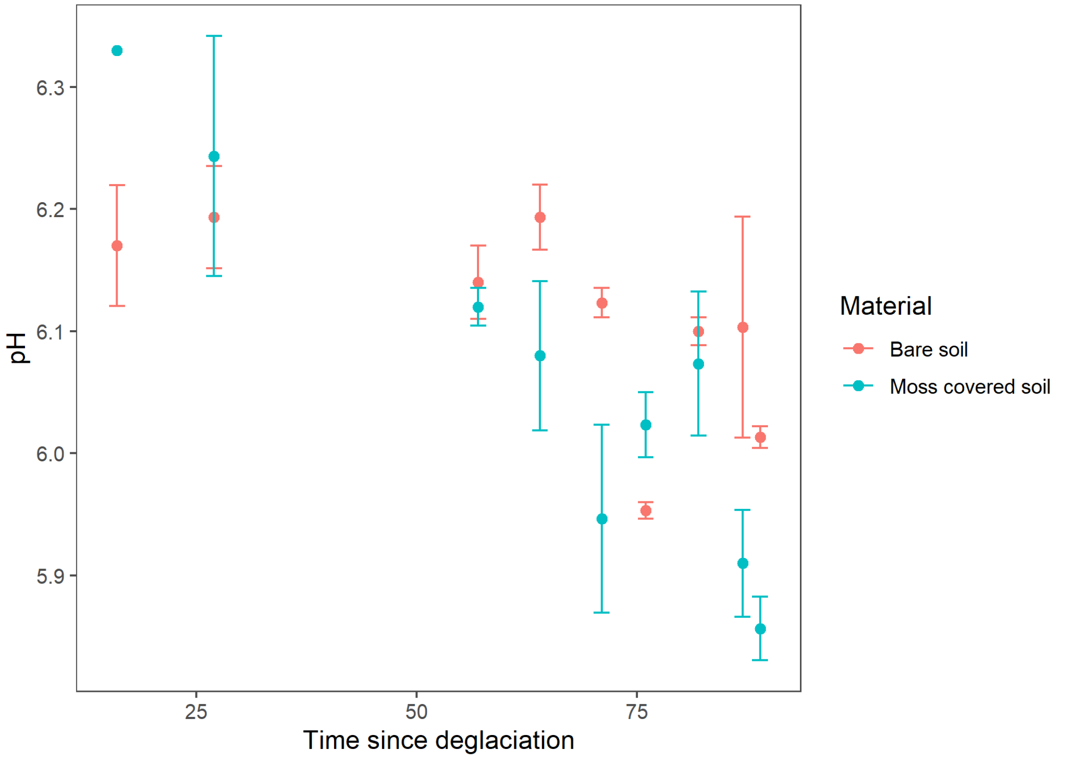


**Table S1.** Correlation matrix of edaphic factors.

| | Time since deglaciation | Species | TC | TN | C:N ratio | Shoot length |
|---|---|---|---|---|---|---|
| Species | 0.862 | | | | | |
| TC | 0.711 | 0.676 | | | | |
| TN | -0.074 | -0.221 | 0.287 | | | |
| C:N ratio | 0.661 | 0.704 | 0.582 | -0.543 | | |
| Shoot length | 0.728 | 0.628 | 0.549 | -0.347 | 0.728 | |
| Moisture content | -0.250 | -0.414 | -0.233 | 0.252 | -0.402 | -0.401 |

**Table S2.** Moss functional traits, moss-associated *nifH* gene abundance and acetylene reduction rates along the chronosequence. For all measured variables mean ± S.E. are shown.

| | Sampling site (time since deglaciation) | | | | | | | | |
|---|---|---|---|---|---|---|---|---|---|
| | 16 | 27 | 57 | 64 | 71 | 76 | 82 | 87 | 89 |
| TC (%) | 21.6 ± 1.8 | 13.8 ± 0.8 | 23.9 ± 4.0 | 24.4 ± 1.9 | 26.4 ± 0.9 | 32.5 ± 3.4 | 28.3 ± 1.9 | 27.7 ± 2.1 | 27.8 ± 2.5 |
| TN (%) | 0.3 ± 0.0 | 0.2 ± 0.0 | 0.3 ± 0.1 | 0.3 ± 0.1 | 0.2 ± 0.0 | 0.2 ± 0.0 | 0.3 ± 0.0 | 0.2 ± 0.1 | 0.2 ± 0.0 |
| C:N ratio | 70.8 ± 7.7 | 67.3 ± 2.0 | 74.8 ± 7.1 | 81.2 ± 12.2 | 127 ± 12.2 | 139.6 ± 5.6 | 112 ± 13.6 | 140 ± 36.7 | 117 ± 6.9 |
| $\delta^{13}C$ (‰) | -28.4 ± 0.1 | -28.1 ± 0.1 | -28.4 ± 0.3 | -26.5 ± 0.2 | -25.7 ± 0.3 | -25.7 ± 0.3 | -25.9 ± 0.3 | -25.8 ± 0.4 | -25.8 ± 0.04 |
| Moisture content (%) | 67.6 ± 8.3 | 71.9 ± 1.5 | 76.3 ± 0.3 | 71.2 ± 1.5 | 58.9 ± 2.1 | 67.4 ± 6.9 | 62.2 ± 2.8 | 59.6 ± 4.5 | 66.1 ± 2.1 |
| Shoot length (mm) | 17.1 ± 5.3 | 11.7 ± 1.7 | 28.1 ± 6.7 | 23.5 ± 0.4 | 42.2 ± 1.0 | 35.8 ± 0.9 | 26.0 ± 2.1 | 37.9 ± 6.5 | 46.9 ± 4.8 |
| *nifH* gene abundance (copies $g^{-1}$) | 184992 ± 85656 | 25421 ± 8832 | 931 ± 318 | 13337 ± 3620 | 19059 ± 8014 | 20439 ± 2083 | 11842 ± 8822 | 6167 ± 1161 | 17269 ± 9588 |
| Acetylene reduction (mol $C^2H^2$ $Kg^{-1}$ $day^{-1}$) | 1.0e-06 ± 2.3e-06 | 4.2e-07 ± 2.6e-07 | 5.7e-06 ± 4.8e-06 | 4.6e-07 ± 4.2e-08 | 3.3e-07 ± 4.8e-08 | 4.7e-07 ± 1.6e-07 | 1.1e-06 ± 5.4e-07 | 6.9e-07 ± 1.4e-07 | 9.1e-07 ± 5.5e-07 |

**Table S3.** Model summary for the linear relationship between shoot length and time since deglaciation in the two moss species. The intercept is for *R. ericoides*.

| term | estimate | std.error | T value | p value |
|---|---|---|---|---|
| Intercept | 30.617 | 5.620 | 5.448 | 1.79e-05 |
| *R. lanuginosum* | -0.002 | 7.807 | 0.000 | 0.999 |
| Time since deglaciation | 9.296 | 3.679 | 2.527 | 0.019 |
| Residual standard error | 9.222 on 22 degrees of freedom | | | |
| Multiple R2 | 0.531 | | | |
| Adjusted R2 | 0.488 | | | |
| F-statistic | 12.44 on 2 and 22 DF | | | |
| p-value | 2.426e-04 | | | |

**Table S4.** Model summary for the linear relationship between moss moisture content and time since deglaciation in the two moss species. The intercept is for *R. ericoides*.

| term | estimate | std.error | T value | p value |
|---|---|---|---|---|
| Intercept | 75.034 | 4.545 | 16.509 | 7.04e-14 |
| *R. lanuginosum* | -13.068 | 6.314 | -2.070 | 0.050 |
| Time since deglaciation | 3.314 | 2.975 | 1.114 | 0.277 |
| Res standard error | 7.459 on 22 degrees of freedom | | | |
| Multiple $R^2$ | 0.215 | | | |
| Adjusted $R^2$ | 0.144 | | | |
| F-statistic | 3.019 on 2 and 22 DF | | | |
| p-value | 0.069 | | | |

**Table S5.** Model summary for the linear relationship between moss TN and time since deglaciation in the two moss species. The intercept is for *R. ericoides*.

| term | estimate | std.error | T value | p value |
|---|---|---|---|---|
| Intercept | 0.317 | 0.046 | 6.932 | 5.84e-07 |
| *R. lanuginosum* | -0.097 | 0.063 | -1.533 | 0.140 |
| Time since deglaciation | 0.034 | 0.030 | 1.136 | 0.268 |
| Res standard error | 0.075 on 22 degrees of freedom | | | |
| Multiple $R^2$ | 0.101 | | | |
| Adjusted $R^2$ | 0.020 | | | |
| F-statistic | 1.242 on 2 and 22 DF | | | |
| p-value | 0.308 | | | |

**Table S6.** Model summary for the linear relationship between moss TC and time since deglaciation in the two moss species. The intercept is for *R. ericoides*.

| term | estimate | std.error | T value | p value |
|---|---|---|---|---|
| Intercept | 23.070 | 2.782 | 8.292 | 3.24e-08 |
| *R. lanuginosum* | 3.284 | 3.865 | 0.850 | 0.405 |
| Time since deglaciation | 3.108 | 1.821 | 1.707 | 0.102 |
| Res standard error | 4.565 on 22 degrees of freedom | | | |
| Multiple $R^2$ | 0.521 | | | |
| Adjusted $R^2$ | 0.477 | | | |
| F-statistic | 11.96 on 2 and 22 DF | | | |
| p-value | 3.052e-04 | | | |

**Table S7.** Model summary for the linear relationship between moss CN ratio and time since deglaciation in the two moss species. The intercept is for *R. ericoides*.

| term | estimate | std.error | T value | p value |
|---|---|---|---|---|
| Intercept | 80.23 | 15.72 | 5.104 | 4.1e-05 |
| *R. lanuginosum* | 38.63 | 21.84 | 1.769 | 0.091 |
| Time since deglaciation | 7.33 | 10.29 | 0.712 | 0.484 |
| Res standard error | 25.8 on 22 degrees of freedom | | | |
| Multiple $R^2$ | 0.507 | | | |
| Adjusted $R^2$ | 0.462 | | | |
| F-statistic | 11.32 on 2 and 22 DF | | | |
| p-value | 4.161e-04 | | | |

**Table S8.** Model summary for the linear relationship between moss $\delta^{13}C$ and time since deglaciation in the two moss species. The intercept is for *R. ericoides*.

| term | estimate | std.error | T value | p value |
|---|---|---|---|---|
| Intercept | -28.282 | 0.256 | -110.522 | < 2e-16 |
| *R. lanuginosum* | 2.442 | 0.356 | 6.870 | 6.7e-07 |
| Time since deglaciation | 0.013 | 0.168 | 0.077 | 0.939 |
| Res standard error | 0.419 on 22 degrees of freedom | | | |
| Multiple $R^2$ | 0.895 | | | |
| Adjusted $R^2$ | 0.886 | | | |
| F-statistic | 93.79 on 2 and 22 DF | | | |
| p-value | 1.705e-11 | | | |

**Table S9.** Model summary for the linear relationship between moss *nifH* gene abundance and time since deglaciation in the two moss species. The intercept is for *R. ericoides*.

| term | estimate | std.error | T value | p value |
|---|---|---|---|---|
| Intercept | -14549 | 36418 | -0.399 | 0.693 |
| *R. lanuginosum* | 67479 | 50590 | 1.334 | 0.196 |
| Time since deglaciation | -64786 | 23839 | -2.718 | 0.013 |
| Res standard error | 59760 on 22 degrees of freedom | | | |
| Multiple $R^2$ | 0.341 | | | |
| Adjusted $R^2$ | 0.281 | | | |
| F-statistic | 5.679 on 2 and 22 DF | | | |
| p-value | 1.02e-02 | | | |

**Table S10.** Model summary for the linear relationship between moss acetylene reduction and time since deglaciation in the two moss species. The intercept is for *R. ericoides*.

| term | estimate | std.error | T value | p value |
|---|---|---|---|---|
| Intercept | 1.018e-06 | 4.015e-07 | 2.536 | 0.020 |
| *R. lanuginosum* | -4.594e-07 | 5.529e-07 | -0.831 | 0.416 |
| Time since deglaciation | 2.126e-07 | 2.474e-07 | 0.859 | 0.400 |
| Res standard error | 5.658e-07 on 20 degrees of freedom | | | |
| Multiple $R^2$ | 0.037 | | | |
| Adjusted $R^2$ | -0.060 | | | |
| F-statistic | 0.381 on 2 and 20 DF | | | |
| p-value | 0.689 | | | |

**Table S11.** Model summary for the linear relationship between ASV richness and time since deglaciation in the two moss species and the soil. The intercept is for *R. ericoides*.

| term | estimate | std.error | T value | p value |
|---|---|---|---|---|
| Intercept | 390.548 | 39.921 | 9.783 | 1.67e-12 |
| *R. lanuginosum* | -2.822 | 50.782 | -0.056 | 0.956 |
| Soil | 159.511 | 45.396 | 3.514 | 0.001 |
| Time since deglaciation | 32.114 | 17.616 | 1.823 | 0.075 |
| Res standard error | 85.82 on 43 degrees of freedom | | | |
| Multiple $R^2$ | 0.547 | | | |
| Adjusted $R^2$ | 0.515 | | | |
| F-statistic | 17.29 on 3 and 43 DF | | | |
| p-value | 1.636e-07 | | | |

**Table S12.** Summary for the post-hoc Tukey test between the ASV richness of two moss species and the soil.

| Comparison | Mean difference | Lower | Upper | P adjusted |
|---|---|---|---|---|
| R. lanuginosum-R. ericoides | 60.985 | -30.584 | 152.555 | 2.499e-01 |
| Soil-R. ericoides | 211.250 | 123.0735 | 152.555 | 1.896e-06 |
| Soil-R. lanuginosum | 150.265 | 81.297 | 219.232 | 1.105e-05 |

**Table S13.** Model summary for the linear relationship between Shannon diversity and time since deglaciation in the two moss species and the soil. The intercept is for *R. ericoides*.

| term | estimate | std.error | T value | p value |
|---|---|---|---|---|
| Intercept | 5.804 | 0.104 | 55.860 | < 2e-16 |
| *R. lanuginosum* | 0.064 | 0.132 | 0.484 | 0.631 |
| Soil | 0.426 | 0.118 | 3.603 | 0.001 |
| Time since deglaciation | 0.068 | 0.046 | 1.491 | 0.143 |
| Res standard error | 0.223 on 43 degrees of freedom | | | |
| Multiple $R^2$ | 0.506 | | | |
| Adjusted $R^2$ | 0.472 | | | |
| F-statistic | 14.7 on 3 and 43 DF | | | |
| p-value | 9.928e-07 | | | |

**Table S14.** Summary for the post-hoc Tukey test between the Shannon diversity of two moss species and the soil.

| Comparison | Mean difference | Lower | Upper | P adjusted |
|---|---|---|---|---|
| R. lanuginosum-R. ericoides | 0.200 | -0.036 | 0.435 | 1.104e-01 |
| Soil-R. ericoides | 0.536 | 0.309 | 0.763 | 2.476e-06 |
| Soil-R. lanuginosum | 0.336 | 0.159 | 0.513 | 1.060e-04 |

**Table S15.** Model summary for the linear relationship between Faith's phylogenetic diversity and time since deglaciation in the two moss species and the soil. The intercept is for *R. ericoides*.

| term | estimate | std.error | T value | p value |
|---|---|---|---|---|
| Intercept | 53.250 | 3.967 | 13.423 | < 2e-16 |
| *R. lanuginosum* | -3.509 | 5.046 | -0.695 | 0.491 |
| Soil | 19.247 | 4.511 | 4.267 | **0.000** |
| Time since deglaciation | 3.105 | 1.750 | 1.774 | 0.083 |
| Res standard error | 8.528 on 43 degrees of freedom | | | |
| Multiple $R^2$ | 0.663 | | | |
| Adjusted $R^2$ | 0.640 | | | |
| F-statistic | 28.24 on 3 and 43 DF | | | |
| p-value | 3.003e-10 | | | |

**Table S16.** Summary for the post-hoc Tukey test between the Faith's phylogenetic diversity of two moss species and the soil.

| Comparison | Mean difference | Lower | Upper | P adjusted |
|---|---|---|---|---|
| R. lanuginosum-R. ericoides | 2.660 | -6.422 | 11.742 | 7.587e-01 |
| Soil-R. ericoides | 24.249 | 15.504 | 32.994 | 8.610e-08 |
| Soil-R. lanuginosum | 21.589 | 14.749 | 28.429 | 3.783e-09 |

**Table S17.** Summary for the Permanova testing the effect of material (moss versus soil) on the bacterial community variation of the mosses and the soil. Here we used time since deglaciation as strata.

| Source | Df | Sum of Squares | Mean Squares | F | $R^2$ | P |
|---|---|---|---|---|---|---|
| Material (moss versus soil) | 1 | 3.405 | 3.405 | 31.371 | 0.411 | **9.999e-05** |
| Residuals | 45 | 4.885 | 0.109 | | 0.589 | |
| Total | 46 | 8.290 | | | 1 | |

**Table S18.** Summary for the Permanova testing the effect of time since deglaciation and moss characteristics on the structure of the soil bacterial community. Here we used moss species as strata.

| Source | Df | Sum of Squares | Mean Squares | F | $R^2$ | P |
|---|---|---|---|---|---|---|
| Time since deglaciation | 1 | 0.009 | 0.009 | 2.444 | 0.104 | **0.042** |
| CN ratio | 1 | 0.007 | 0.008 | 2.01 | 0.086 | **0.044** |
| TN | 1 | 0.002 | 0.002 | 0.57 | 0.024 | 0.758 |
| Moisture content | 1 | 0.005 | 0.005 | 1.418 | 0.060 | 0.176 |
| Residuals | 17 | 0.066 | 0.004 | | 0.725 | |
| Total | 21 | 0.091 | | | 1 | |

**Table S19.** Summary for the Permanova testing the effect of time since deglaciation and moss characteristics on the structure of the moss bacterial community variation. Note that we used moss species as strata here.

| Source | Df | Sum of Squares | Mean Squares | F | $R^2$ | P |
|---|---|---|---|---|---|---|
| Time since deglaciation | 1 | 0.099 | 0.099 | 18.230 | 0.381 | **9.99e-05** |
| CN ratio | 1 | 0.009 | 0.009 | 1.662 | 0.035 | 0.214 |
| TN | 1 | 0.011 | 0.011 | 1.980 | 0.041 | 0.065 |
| Moisture content | 1 | 0.033 | 0.033 | 5.939 | 0.124 | **9.999e-05** |
| Residuals | 20 | 0.110 | 0.005 | | 0.418 | |
| Total | 24 | 0.262 | | | 1 | |

**Table S20.** Summary for the Permanova testing the effect of time since deglaciation and moss characteristics on the structure of the bacterial community of the moss *R. ericoides*.

| Source | Df | Sum of Squares | Mean Squares | F | $R^2$ | P |
|---|---|---|---|---|---|---|
| Time since deglaciation | 1 | 0.006 | 0.006 | 2.876 | 0.216 | **0.004** |
| CN ratio | 1 | 0.005 | 0.005 | 2.491 | 0.188 | **0.022** |
| TN | 1 | 0.003 | 0.003 | 1.787 | 0.134 | 0.093 |
| Moisture content | 1 | 0.006 | 0.006 | 3.134 | 0.236 | **0.004** |
| Residuals | 3 | 0.006 | 0.002 | | 0.226 | |
| Total | 7 | 0.026 | | | 1 | |

**Table S21.** Summary for the Permanova testing the effect of time since deglaciation and moss characteristics on the structure of the bacterial community of the moss *R. lanuginosum*.

| Source | Df | Sum of Squares | Mean Squares | F | $R^2$ | P |
|---|---|---|---|---|---|---|
| Time since deglaciation | 1 | 0.005 | 0.005 | 1.923 | 0.094 | 0.079 |
| CN ratio | 1 | 0.004 | 0.004 | 1.658 | 0.081 | 0.128 |
| TN | 1 | 0.003 | 0.003 | 1.123 | 0.055 | 0.330 |
| Moisture content | 1 | 0.009 | 0.009 | 3.844 | 0.187 | **0.003** |
| Residuals | 12 | 0.029 | 0.002 | | 0.584 | |
| Total | 16 | 0.050 | | | 1 | |

**Table S22.** Statistics of the structural equation model of direct and indirect effects of warming on $N_2$-fixation as shown in Figure 7. We show the standardized path coefficients (Std. est.), the standard error of regression weight (se), the z-value (z) and the significance level for the regression weight (p). Significant effects are shown in bold.

| Parameter | Variable | Std. est. | se | z | P-value |
|---|---|---|---|---|---|
| Moss bacterial community | Time since deglaciation (tm) | -0.33 | 0.17 | -1.88 | 0.06 |
| | TN (Nm) | 0.41 | 0.16 | 2.55 | **0.01** |
| | Moisture (Mm) | -0.63 | 0.16 | -4.01 | **6.21E-05** |
| *nifH* | axis1 (mn) | -0.87 | 0.22 | -3.97 | **7.18E-05** |
| | Time since deglaciation (tn) | -0.28 | 0.20 | -1.38 | 0.17 |
| | TN (Nn) | 0.05 | 0.20 | 0.23 | 0.82 |
| | Moisture (Mn) | -0.23 | 0.25 | -0.92 | 0.36 |
| $N_2$-fixation | axis1 (mnfix) | -0.63 | 0.39 | -1.61 | 0.11 |
| | nifH (nnfix) | -0.68 | 0.27 | -2.54 | **0.01** |
| | Time since deglaciation (tnfix) | 0.11 | 0.25 | 0.42 | 0.67 |
| | TN (Nnfix) | 0.11 | 0.24 | 0.45 | 0.65 |
| | Moisture (Mnfix) | -0.27 | 0.29 | -0.92 | 0.36 |
| TN | Time since deglaciation (tC) | -0.13 | 0.25 | -0.52 | 0.61 |
| Moisture | Time since deglaciation (tM) | -0.30 | 0.23 | -1.31 | 0.19 |
| Indirect effects on $N_2$-fixation | | Std. est. | se | z | P-value |
| Total_effect_nfix | tnfix+(tM*Mnfix)+(tM*Mn*nnfix)+(tM*Mm*mnfix)+(tM*Mm*mn*nnfix)+(tN*Nnfix)+(tN*Nn*nnfix)+(tN*Nm*mnfix)+(tN*Nm*mn*nnfix)+(tm*mnfix)+(tm*mn*nnfix)+(tn*nnfix) | 0.33 | 0.22 | 1.47 | 0.14 |
| Indirect_time_nfix | (tM*Mnfix)+(tM*Mn*nnfix)+(tM*Mm*mnfix)+(tM*Mm*mn*nnfix)+(tN*Nnfix)+(tN*Nn*nnfix)+(tN*Nm*mnfix)+(tN*Nm*mn*nnfix)+(tm*mnfix)+(tm*mn*nnfix)+(tn*nnfix) | 0.22 | 0.18 | 1.22 | 0.22 |
| Indirect_time_moisturen_nfixall | (tM*Mnfix)+(tM*Mn*nnfix)+(tM*Mm*mnfix)+(tM*Mm*mn*nnfix) | 0.03 | 0.08 | 0.38 | 0.71 |

| | | | | | |
|---|---|---|---|---|---|
| Indirect_time_ moisturen_nfix 1 | (tM*Mnfix) | 0.08 | 0.11 | 0.75 | 0.46 |
| Indirect_time_ moisturen_nfix 2 | (tM*Mn*nnfix) | -0.05 | 0.07 | -0.71 | 0.48 |
| Indirect_time_ moisturen_nfix 3 | (tM*Mm*mnfix) | -0.12 | 0.12 | -0.96 | 0.34 |
| Indirect_time_ moisturen_nfix 4 | (tM*Mm*mn*nnfix) | 0.11 | 0.11 | 1.03 | 0.31 |
| Indirect_time_ TN_nfixall | (tN*Nnfix)+(tN*Nn*nnfix)+(tN*Nm*mnfix)+(tN* Nm*mn*nnfix) | -0.00 | 0.03 | -0.24 | 0.81 |
| Indirect_time_ TN_nfix1 | (tN*Nnfix) | -0.01 | 0.04 | -0.34 | 0.74 |
| Indirect_time_ TN_nfix2 | (tN*Nn*nnfix) | 0.00 | 0.02 | 0.21 | 0.84 |
| Indirect_time_ TN_nfix3 | (tN*Nm*mnfix) | 0.03 | 0.07 | 0.48 | 0.63 |
| Indirect_time_ TN_nfix4 | (tN*Nm*mn*nnfix) | -0.03 | 0.06 | -0.48 | 0.63 |
| Indirect_micro b_nfix1 | (Nm*mn*nnfix) | 0.25 | 0.16 | 1.55 | 0.12 |
| Indirect_micro b_nfix2 | (mn*nnfix) | 0.60 | 0.30 | 1.99 | **0.05** |
| Indirect_micro b_nfix3 | (Mm*mn*nnfix) | -0.38 | 0.22 | -1.74 | 0.08 |
| Indirect_time_ microb_nfixall | (tm*mnfix)+(tm*mn*nnfix) | 0.01 | 0.12 | 0.09 | 0.93 |
| Indirect_time_ microb_nfix1 | (tm*mnfix) | 0.21 | 0.17 | 1.23 | 0.22 |
| Indirect_time_ microb_nfix2 | (tm*mn*nnfix) | -0.20 | 0.14 | -1.38 | 0.17 |
| Indirect_time_ nifh_nfix | (tn*nnfix) | 0.19 | 0.15 | 1.25 | 0.21 |

**Table S23.** Model summary for the linear relationship between soil moisture and time since deglaciation in bare and moss-covered soil. The intercept is for *bare soil.*

| term | estimate | std.error | T value | p value |
|---|---|---|---|---|
| Intercept | 2.394 | 0.074 | 32.529 | < 2e-16 |
| Time since deglaciation | -0.002 | 0.001 | -2.398 | < 0.05 |
| Moss-covered soil | 0.198 | 0.050 | 3.928 | < 0.001 |
| Res standard error | 0.184 on 50 degrees of freedom | | | |
| Multiple $R^2$ | 0.296 | | | |
| Adjusted $R^2$ | 0.268 | | | |
| F-statistic | 10.49 on 2 and 50 DF | | | |
| p-value | < 0.001 | | | |

**Table S24.** Model summary for the linear relationship between soil pH and time since deglaciation in bare and moss-covered soil. The intercept is for *bare soil.*

| term | estimate | std.error | T value | p value |
|---|---|---|---|---|
| Intercept | 1.845e+00 | 7.051e-03 | 261.692 | < 2e-16 |
| Time since deglaciation | -5.605e-04 | 9.956e-05 | -5.630 | < 0.001 |
| Moss-covered soil | -8.999e-03 | 4.599e-03 | -1.957 | 0.0561 |
| Res standard error | 0.017on 49 degrees of freedom | | | |
| Multiple $R^2$ | 0.434 | | | |
| Adjusted $R^2$ | 0.411 | | | |
| F-statistic | 18.79 on 2 and 49 DF | | | |
| p-value | 8.773e-07 | | | |

**Supplementary Methods 1** Soil parameters of bare and moss-covered soils in the glacier forefields

Samples were collected in late April 2021 along the same transect as the previous sample collection in 2018 had taken place. The same coordinates were visited. At each sampling point, three bare soil samples and three moss-covered soil samples were taken, both the upper 10 cm of the soil, but without the moss mat.

The soil samples were stored cool until processed. Field-moist soil samples were sieved to 2 mm. For soil moisture content, samples were weighed before and after drying at 70 °C for 24 h. pH was measured after mixing 5 g of soil and 15 ml deionized water for 1 h and left to stand overnight.

One soil moisture measurement was left out as the weight after drying was registered as higher than before drying. And two pH measurements were ignored as the pH meter was not calibrated and the samples were discarded.

Multiple linear regressions with time since deglaciation and material (bare soil versus moss-covered soil) as independent variables were used to test whether soil moisture, pH and organic matter content change with time since deglaciation and differ between material.
